# Asymmetric adhesion of rod-shaped bacteria controls microcolony morphogenesis

**DOI:** 10.1101/104679

**Authors:** Duvernoy Marie-Cécilia, Mora Thierry, Ardré Maxime, Croquette Vincent, Bensimon David, Quilliet Catherine, Ghigo Jean-Marc, Balland Martial, Beloin Christophe, Lecuyer Sigolène, Desprat Nicolas

## Abstract

Bacterial biofilms are spatially structured communities, within which bacteria can differentiate depending on environmental conditions. During biofilm formation, bacteria attach to a surface and use cell-cell contacts to convey the signals required for the coordination of biofilm morphogenesis. How bacteria can maintain both substrate adhesions and cell-cell contacts during the expansion of a microcolony is still a critical yet poorly understood phenomenon. Here, we describe the development of time-resolved methods to measure substrate adhesion at the single cell level during the formation of *E. coli* and *P. aeruginosa* microcolonies. We show that bacterial adhesion is asymmetrically distributed along the cell body. Higher adhesion forces at old poles put the daughter cells under tension and force them to slide along each other. These rearrangements increase cell-cell contacts and the circularity of the colony. We propose a mechanical model based on the microscopic details of adhesive links, which recapitulates microcolony morphogenesis and quantitatively predicts bacterial adhesion from simple time lapse movies. These results explain how the distribution of adhesion forces at the subcellular level directs the shape of bacterial colonies, which ultimately dictates the circulation of secreted signals.

---

From microbiome to nosocomial diseases, commensal and pathogenic biofilms have a crucial impact on human lifestyle. Attached to biotic or synthetic surfaces, biofilms form reproducible and organized structures [1, 2] that respond to environmental conditions [3, 4]. Inside biofilms, the diffusion and consumption of metabolites generate spatial heterogeneities [5, 6, 7] that further increase the palette of group strategies [8, 9, 10, 11, 12, 13]. Signals that regulate bacterial behaviors during colony morphogenesis not only diffuse through an amorphous mass of bacteria, but are also mediated by cell-cell contacts. Direct contacts serve as cues for delimiting the boundaries of the colony exposed to non-kin bacteria [14, 15], coordinate cell movements [16, 17], convey signals through the colony [18] and promote the use of public good secretion [19]. However, bacterial elongation, which drives the expansion of the microcolony, competes with both cell-cell cohesion and surface attachment. To understand how elongation and adhesion shape morphogenesis, we performed a large set of experiments at both cellular and microcolony scales.

Heterogeneous elongation and/or differential adhesion can shape bacterial microcolonies. We first investigated the contribution of elongation by tracking individual cells within WT microcolonies of *E. coli* and *P. aeruginosa* (Figure S1A) growing between a soft agarose gel (1%) and a glass coverslip (Fig. 1A, Movie S1). Surprisingly, bacteria elongate at the same rate regardless of their position within the microcolony (Figure S1B). As a result bacteria are pushed by others during the expansion of the microcolony on the surface (Fig. 1C,D and Figure S1C). Bacteria at the periphery of the microcolony, which are pushed by more siblings, experience larger displacements (Figure S2A,B) and the oldest cells [20] are systematically located at the periphery of the microcolony (Fig. 1E and Figure S1D). On average, bacteria are pushed radially and outwards from the center of the colony (Figure S1C). The cylindrical shape of bacteria constrains their orientation and neighboring bacteria tend to align. For *E. coli* cells whose length and width after division are respectively 4.4µm and 1.4µm, we measured that the 2D nematic order parameter decays exponentially with a characteristic length of 10µm (Figure S2C,D). For *P. aeruginosa* cells, whose length and width are respectively 2.3µm and 0.9µm, the 2D nematic order parameter decays with a characteristic length of 3µm (Figure S2C,D). These results show that cell elongation uniformly drives the expansion of the microcolony. However, the bacteria tend to align and steric interactions participate in shaping the microcolony.

**Figure 1.**
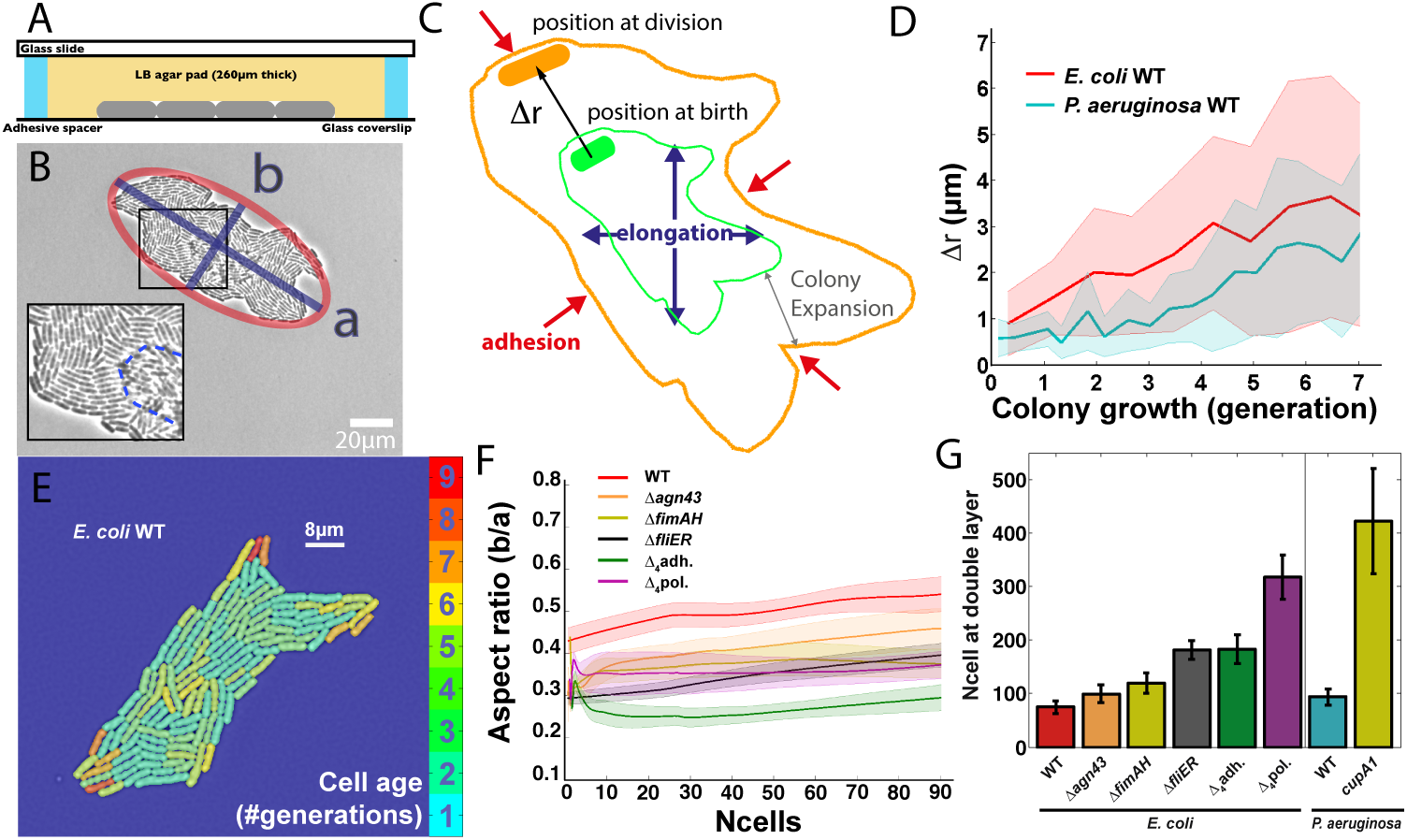
Microcolony morphogenesis.

A. Schematic of the culture chamber. Bacteria are sandwiched between an agarose pad and a glass coverslip. B. Image of a wild type *E. coli* microcolony after the formation of a second layer. The inset highlights the double layer at the center and on top of the microcolony. By image analysis, we compute the long axis *a* and the short axis *b* of the ellipse that fits the mask of the microcolony. C. Mechanisms of microcolony expansion and schematic of cell movements (∆*r*) occurring during each cell cycle. D. Average cell displacement ∆*r* as a function of time from the beginning of microcolony growth for WT *E. coli* and for WT *P. aeruginosa* PAO1. ∆*r* of a given cell is computed as the distance travelled between birth and cell division. E. Distribution of cell age within a WT *E. coli* microcolony. The colors code for the age of bacteria given by the history of their oldest pole (blu is young; red is old). F. Aspect ratio (b/a) for WT *E. coli* and mutants impaired in adhesive properties as a function of the number of bacteria in the microcolony. G. Microcolony size at the onset of the double layer formation for the same set of strains as in F. In panels F, G, error bars represent the standard errors (WT *E. coli*, red, N=7; ∆*agn43* orange, N=5; ∆*fimAH* khaki, N=15; ∆*fliER* black, N=19; Δ_4_pol purple, N=5; Δ_4_adh green, N=5). In panel D, error bars represent the standard deviations (WT *E. coli* red, N=7; WT *P. aeruginosa* cyan, N=10).

We then investigated how adhesion to the substrate constrains the morphogenesis of microcolonies. We quantified the shape of *E. coli* microcolonies by measuring their aspect ratio (Fig. 1B). Using different mutants with impaired adhesion properties [autotransporter Antigen43 (Δ*agn43*), the flagellum (Δ*fliER*), various surface appendages (Δ_4_*adh*; absence of flagella, Ag43, type 1 fimbriae and curli) or exopolysaccharides (Δ_4_*pol*; absence of Yjb polysaccharide, cellulose, PGA and colanic acid)], we show that reducing the level of adhesion gives rise to more elongated microcolonies (Fig. 1F, Movie S2). After a phase of growth in 2D, microcolonies undergo a transition and initiate 3D expansion by forming a second layer at the center of the colony [21, 22] (Fig. 1B inset). This transition also depends on the level of substrate adhesion, since mutants form a double layer at larger microcolony sizes for both *E. coli* and *P. aeruginosa* (Fig. 1G). Thus, adhesion controls both the shape of the microcolony as well as the transition to 3D growth.

Cell elongation and steric repulsion cause collective rearrangements in the colony, which hide potential effects of differential adhesion. To decouple these effects we probed the adhesive properties of single isolated bacteria. We tracked the movement of the center of mass (CM) during the cell cycle. Cell elongation is known to be uniform since cell wall insertion is patchy but homogeneous along the cell axis for many rod-shaped bacteria [23, 24, 25, 26, 27]. We thus fitted the relationship ΔX(*t*) = *A_cell_*Δ*L*(*t*) between the displacement of the CM, ∆*X*, and the change in cell length, ∆*L*, to infer *A*_cell_ a measure of the asymmetry in cell adhesion (Fig. 2A). If adhesion is uniform, isolated bacteria should elongate symmetrically around their center of mass, yielding a null asymmetry parameter (*A_cell_* = 0). On the contrary, we noticed that the CM moves during cell elongation (Movie S3), which indicates that one pole adheres more strongly than the other (Figure S3A,B). In order to test whether the CM displacement is in fact caused by non-uniform adhesion, we compared the distribution of *A_cell_* in populations of WT bacteria against various mutants of adhesion (Fig. 2B). For *E. coli*, the level of asymmetry is reduced by 40% in single adhesin mutants (Δ*agn43*,Δ*fliER*,Δ*fimAH*) and by up to 70% in quadruple mutants (Δ_4_*adh*, Δ_4_pol). This is consistent with the fact that some adhesive factors such as flagella [28], polysaccharide [29], cellulose [30] or the Ag43 adhesin [31] have been shown to be polar. We further analyzed the localization of Ag43 at the single cell level by performing immunofluorescence microscopy on *E. coli* cells. Strikingly, Ag43 is distributed at the surface of single cells with a strong monopolar signal (Figure S3C). Finally, we also observe in *P. aeruginosa* that the level of asymmetry is reduced by 17% in a *cupA1* fimbriae mutant compared to the WT strain (Fig. 2B). These observations suggest that polar adhesion might be a general feature of gram-negative bacteria.

**Figure 2.**
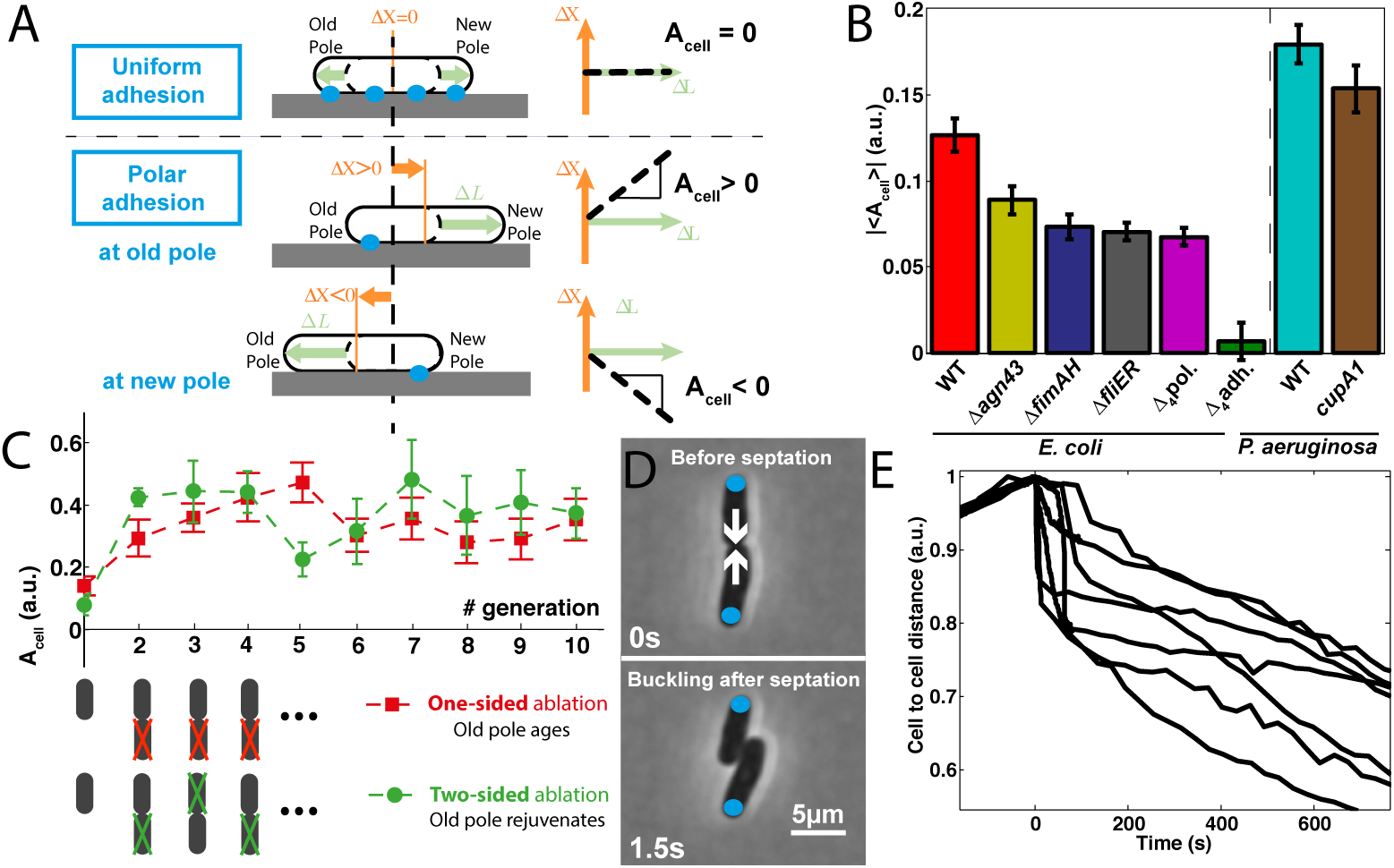
Polar adhesion induces reorganization after the first division.

A. Polar adhesion translates into a displacement of the center of mass (CM) of a growing cell. The level of asymmetry for a cell, *A*_cell_, is defined according to Δ*X* = *A*_cell_ Δ*L*, where Δ*X* is the displacement of the CM projected along the long axis of the cell and Δ*L* is the cell elongation. The cell axis is oriented from the old pole towards the new pole. The asymmetry |*A*_*cell*_| can vary from 0 (uniform adhesion) to 0.5 (polar adhesion). B. Average asymmetry |〈*A*_*cell*_〉| in populations of isolated bacteria for different strains. Since pole history is not known before the first division, we measured the absolute value of the asymmetry on populations of cells: WT *E. coli* (red, N=146) and different mutants deleted for one adhesin factors (∆*agn43* orange, N=116; ∆*fimAH* khaki, N=100; ∆*fliER* black, N=96) or four (Δ_4_*adh* green, N=95; Δ_4_pol purple, N=159) adhesin factors; WT *P. aeruginosa* PA14 (cyan, N=111) and the fimbriae mutant *cupA1* (brown, N=98). Error bars represent standard errors. C. Successive laser ablations of one daughter after division of WT *E. coli* enables to orient the cell axis and assess the sign of *A*_cell_. Positive *A*_cell_ indicates that adhesion is biased towards the old pole. D. Images of WT *E. coli* before and after reorganization. Old pole are marked with a blue dot. E. Distance between the CM of the two daughter cells normalized by the size at division. Before division, the cell to cell distance is computed as half the total cell length. Traces show individual reorganizations. Time is set to 0 when the cell to cell distance is maximal.

To determine which pole carries most of the adhesion, we tracked single isolated cells and performed laser ablation on one of the two daughter cells after each division (Fig. 2C, Movie S4). In a first series of experiments, laser ablations were always performed on the same side of the cell pair, so that the same old pole was conserved throughout the experiment (one-sided ablations, Fig. 2C and Figure S4A,B). Consistent with what we found for a WT population of *E. coli* (Fig. 2B), the absolute value of the asymmetry for the first generation is 0.13 (Fig. 2C). For subsequent generations, the signed asymmetry *A*_cell_ takes positive values, showing that adhesion is stronger at the old pole than at the new pole (Fig. 2C). To rule out possible artifacts due to systematic one-sided ablations, we performed two-sided ablation experiments by targeting the opposite side of the cell pair at each division (Movie S4). In this configuration, the old pole is renewed at each generation (Figure S4A), but A_cell_ remains positive with values similar to the ones obtained for one-sided ablations (Fig. 2C). These experiments show that adhesion at the old pole fully matures over one cell cycle since both modes of ablation yield the same values of the asymmetry.

At division, the old poles of the two daughter bacteria are located on opposite sides of the cell pair (Fig. 2D). Since bacteria are more strongly attached at their old pole, they will tend to elongate towards each other. This situation favors a buckling instability [32, 33] that triggers the rapid reorganization of the two daughter bacteria (Fig. 2E, Movie S5). For WT *E. coli*, we observed that the magnitude of this reorganization is positively correlated to the level of asymmetry of the mother cell (Figure S3D).

This result illustrates how polar adhesion coupled with bacterial elongation can generate mechanical stress. More generally, adherent bacteria that are pushed by neighboring cells should transmit forces to the underlying substrate. Using time-resolved force microscopy [34], we measured the dynamics of the force pattern during the growth of *E. coli* and *P. aeruginosa* microcolonies. Briefly, bacteria were confined between a rigid agarose gel (2%) and a polyacrylamide (PAA) gel, in which embedded fluorescent beads serve as deformation markers. Using inverse Fourier transform, we calculated the stress tensor on a lattice under the colony from the beads displacement field. The mechanical stress bacteria exert on the substrate is heterogeneous and dynamical (Fig. 3A, Figure S5A and Movie S6). Yet, the average radial stress does not depend on the position inside the colony (Figure S5B). The sum of the magnitudes of forces developed by the microcolony on the substrate *F*_colo_ increases linearly with the area of the microcolony, so that the average stress *σ_colo_* is constant during growth (Fig. 3B). The average stress *σ_colo_* is lower for WT *P. aeruginosa* (80Pa) than for WT *E. coli* (150Pa). For both species, adhesion mutants display lower *σ_colo_* (Fig. 3C). We then measured the maximal force *F*_max_ generated at adhesion foci throughout the growth of the microcolony. *F*_max_ increases during early microcolony growth, then saturates (Fig. 3D). We quantified the dynamics of the saturation by fitting our experimental data with an exponential relaxation *F_max_*(*t*) = *F_sat_*[1 − *exp*(−*A*(*t*)/*A*_0_)], where *F_sat_* is the value of the plateau and *A*_0_ the characteristic area at which the curve plateaus. Remarkably, for both species, the saturation is reached for a colony size of about 8 bacteria, corresponding to one cell being fully surrounded by neighbors [35]. The value of the plateau is consistent with previous AFM measurements [36, 37]. The level of saturation *F_sat_* is lower for WT *P. aeruginosa* (57±4pN) than for WT *E. coli* (114±4pN) (Fig. 3E). For *E. coli*, *F_sat_* is reduced by 4% in a mutant of adhesins Δ_4_adh and by 38% in a mutant of polysaccharides Δ_4_pol. For *P. aeruginosa*, the level of saturation is reduced by 23% in the fimbriae mutant PA14*cupA1*. In summary, the average stress over the whole microcolony remains constant during growth. However local fluctuations are present with an amplitude that is bounded by the strength of adhesion.

**Figure 3.**
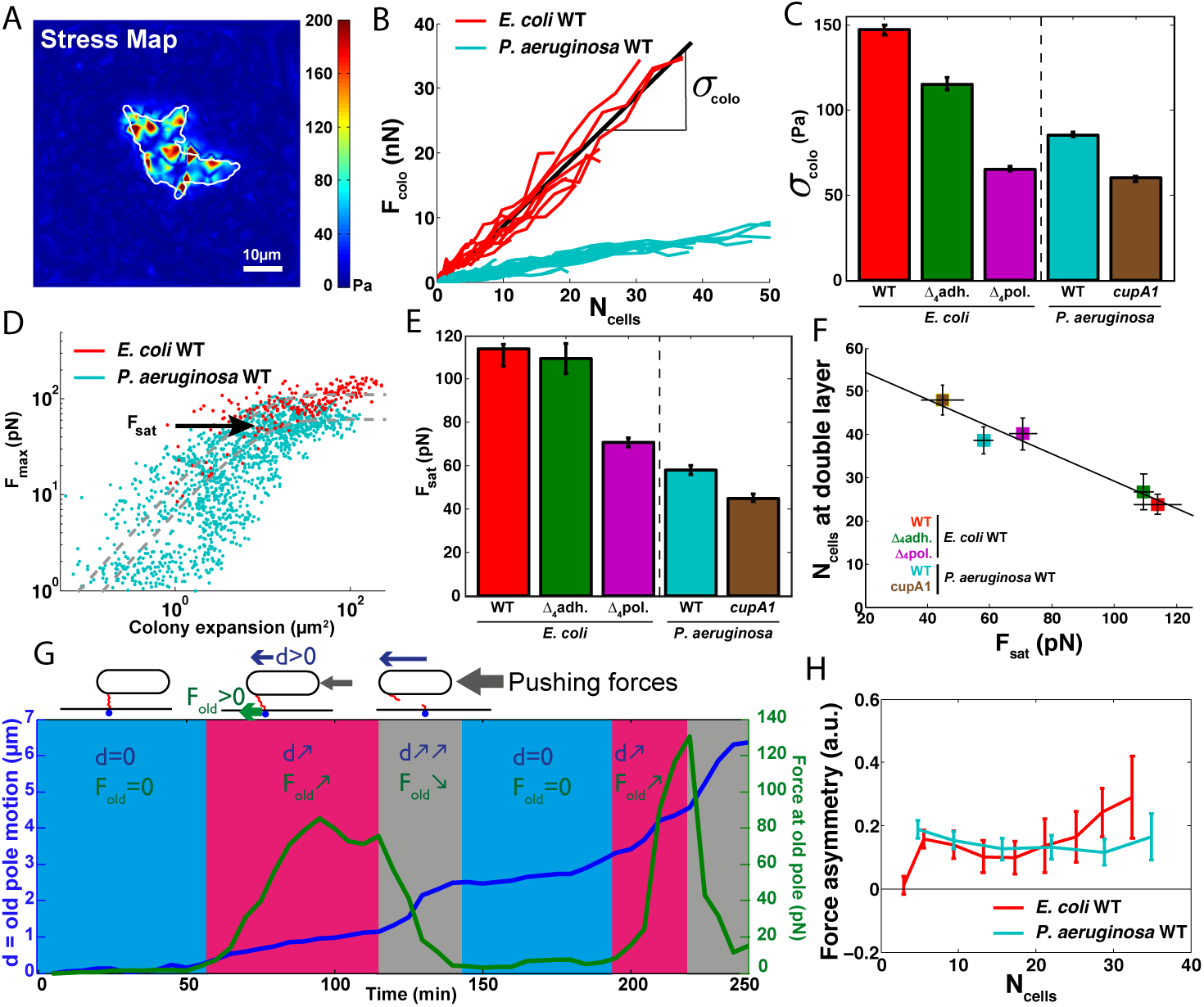
Cell-substrate adhesion dynamics inside the microcolony.

A. Map of the stress that bacteria exert on the substrate at a given time for a growing *E. coli* wild type microcolony (white contour). B. The total force *F*_colo_ developed by independent microcolonies on the substrate increases linearly with the area of the microcolony for both WT *E. coli* (red) and WT *P. aeruginosa* (cyan). We compute the stress *σ_colo_* beneath the microcolony as the average slope of these curves. C. *σ_colo_* for WT *E. coli* (red, N=12) and mutants (Δ_4_adh, green, N=19; Δ_4_pol, purple, N=48); WT *P. aeruginosa* PA14 (cyan, N=16) and the fimbriae mutant *cupA1* (brown, N=14). D. The maximal force measured locally (i.e. for a lattice of characteristic size 510 nm) on the substrate saturates with the microcolony area for WT *E. coli* (red) and WT *P. aeruginosa* (cyan). The saturation force F_sat_ corresponds to the value of the plateau. E. *F_sat_* for the same set of strains displayed in panel C. F. Number of bacteria inside the microcolony at the onset of a second layer as a function of the saturation force for the strains displayed in panel C and E. G. Dynamics of the adhesive force (green) and displacement (blue) at an individual pole. If the pole slightly moves, almost no force is exerted on the substrate (blue phase). The force increases when the pole displaces (pink phase). The force then drops following an abrupt displacement (grey phase). H. Force asymmetry of the microcolony *A*_colo_ for wild type *E. coli* (red, N=12) and *P. aeruginosa* (cyan, N=16) measured as *A_colo_* = [*max*(*F_old_*) − *max*(*F_new_*)]/*F_sat_*. In panel C and E, error bars represent the confidence interval of fitted parameters. In panel H, error bars represent the standard errors.

The saturation of the maximal force suggests that adhesive bonds are broken by the elongation forces of surrounding bacteria. To test this hypothesis, we tracked individual poles during microcolony growth. We measured that the adhesive force at a given pole drops when the pole undergoes a sudden displacement (Fig. 3G). The statistical analysis of pole movements and forces over many microcolonies shows a negative correlation between pole displacement and force variation, indicating that adhesive bonds break under the load generated during the growth of the microcolony (Figure S6A). The saturation level *F_sat_* thus reflects the strength of adhesion for a given strain. Because most of the bacteria are constantly pushed by the elongation of others, not all poles are maintaining their adhesion with the substrate. As a result, non-adhering bacteria slide on the substrate and not all the beads located beneath the microcolony moves (Figure S6B,C). While the ruptures of individual adhesive bonds enable the expansion of the microcolony on the substrate, the formation of a double layer at the center of the microcolony (Fig. 1B inset) [21, 22] suggests that there is a critical colony size at which the force generated by bacterial elongation cannot overcome the cumulated adhesion of surrounding bacteria. Indeed, our experimental measurements do show that the strength of adhesion to the substrate, *F_sat_*, directly sets the number of bacteria in the microcolony at the onset of the second layer formation (Fig. 3F). Finally we investigated if the asymmetry between poles persists at the scale of the microcolony by comparing the forces at new and old poles. In agreement with experiments on individual bacteria, we observed that adhesive forces are stronger at old poles during the growth of both *E. coli* and *P. aeruginosa* microcolonies (Fig. 3H).

Since the temperature modulates both bacterial elongation and protein expression levels, we investigated its role in the adhesive properties of the cells. We compared the set of previous experiments at 34°C with the same ones carried out at 28°C. Lowering the temperature reduces the asymmetry of the WT E. coli cells by 7%, while the asymmetry of the *E. coli* strain deleted for four adhesins Δ_4_adh remains very weak (Figure S7A). Concomitantly, the average stress *σ_colo_*, and saturation force *F_sat_* both decrease with the temperature. For both strains, *σ_colo_* and *F_sat_* drop by respectively 40% and 25% between 34°C and 28°C (Figure S7B, C). Changing the temperature also allowed us to test the mutant strain, *ompR234*, which overexpresses the curli adhesin at 28°C [38]. In the *ompR234* strain, the asymmetry at the single cell level raises by 25% (Figure S8A) and *σ_colo_* increases by 15% compared to the WT *E. coli* strain (Figure S8B). However, *F_sat_* only increases by 7% (Figure S8C). This weaker effect of curli over-expression on the saturation force might indicate that the value of the saturation force does not simply scale with the expression rate. Similarly to experiments at 34°C, the number of bacteria in the microcolony at the onset of the double layer formation linearly decreases with the strength of cell-substrate adhesion at 28°C (Figure S8D). However, the slope of the relationship is 4 times steeper at 28°C than at 34°C (Figure S7D). If raising the temperature increases both the adhesive properties and the proliferation of cells, it modifies the transition to 3D in a non trivial manner.

To get a comprehensive view of microcolony morphogenesis, we propose a computational model that integrates our observations. Bacteria are simulated as 2D spherocylinders that interact with neighbouring cells via a short range attractive force and an elastic repulsion force. Each cell interacts with the substrate by means of adhesive links that form at a fixed rate at both poles. New poles are thus free of adhesive molecules right after septation and the asymmetry is a natural consequence of cell division. Adhesive links are ruptured once the force they experience reaches a threshold *F_link_*, which directly sets the saturation force *F_sat_*. For a rupture force *F_link_* of 4.25pN that yields the experimentally measured *F_sat_* = 115pN for *E. coli*, our simulation predicts the shape of the microcolony in excellent agreement with experimental results (Fig. 4A) and also reproduce the dynamical heterogeneity observed on the stress map (Movie S7). Consistent with experiments, a weaker adhesive force on individual links generates more elongated microcolonies (Figure S9A,B,C). To understand the transition from a 2D growth to a 3D expansion, we investigated how the elastic energy *E_adh_* stored in the adhesive links and the repulsive energy *E_rep_* between bacteria scale with scale with the size of the microcolony for different values of the saturation force *F_sat_*. We found that 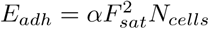 and 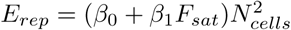 Fig. 4C,D). Thanks to these scalings, we derived the analytical expression (see Methods) for the 2D-3D transition by comparing the situation in which the microcolony expands entirely in the plane to a situation in which one bacterium deforms the gel above to expand into the third dimension (Fig. 4B and Figure S10). With the Young modulus of the PAA gel, this analytical expression fits the 3D transition behavior in the force measurement experiments (Fig. 4E inset). To go beyond force measurement experiments, we tried to infer the adhesion forces in the agarose-glass experiments where they can not directly be measured. Keeping the same equation (see Methods) but using the Young modulus of the agarose gel (Fig. 4E), we deduced the saturation force *F_sat_* corresponding to the number of bacteria measured at the onset of the second layer (Fig. 1G). We then inverse the relation between *F_sat_* and *F_link_* (Figure S9D). Finally, we were able to reproduce the experimental shape and organization of microcolonies in agarose-glass configuration (Fig. 4F,G) simply by using *F_link_* deduced from an easily measurable parameter. To summarize our results, microcolonies grown between agarose and PAA are more elongated than between agarose and glass due to a weaker adhesion. In addition, they develop a double layer earlier since the PAA gel is softer (4kPa) than the agarose (15kPa). We further addressed the role of polar adhesion by performing simulations where adhesive links are allowed to form homogeneously along the cell body. We show that for the same set of parameters, uniform adhesion leads to more elongated colonies in which bacteria are more aligned (Fig. 4F,G). Hence, polar adhesion decreases the orientational order, which subsequently enables the formation of rounder microcolonies.

**Figure 4.**
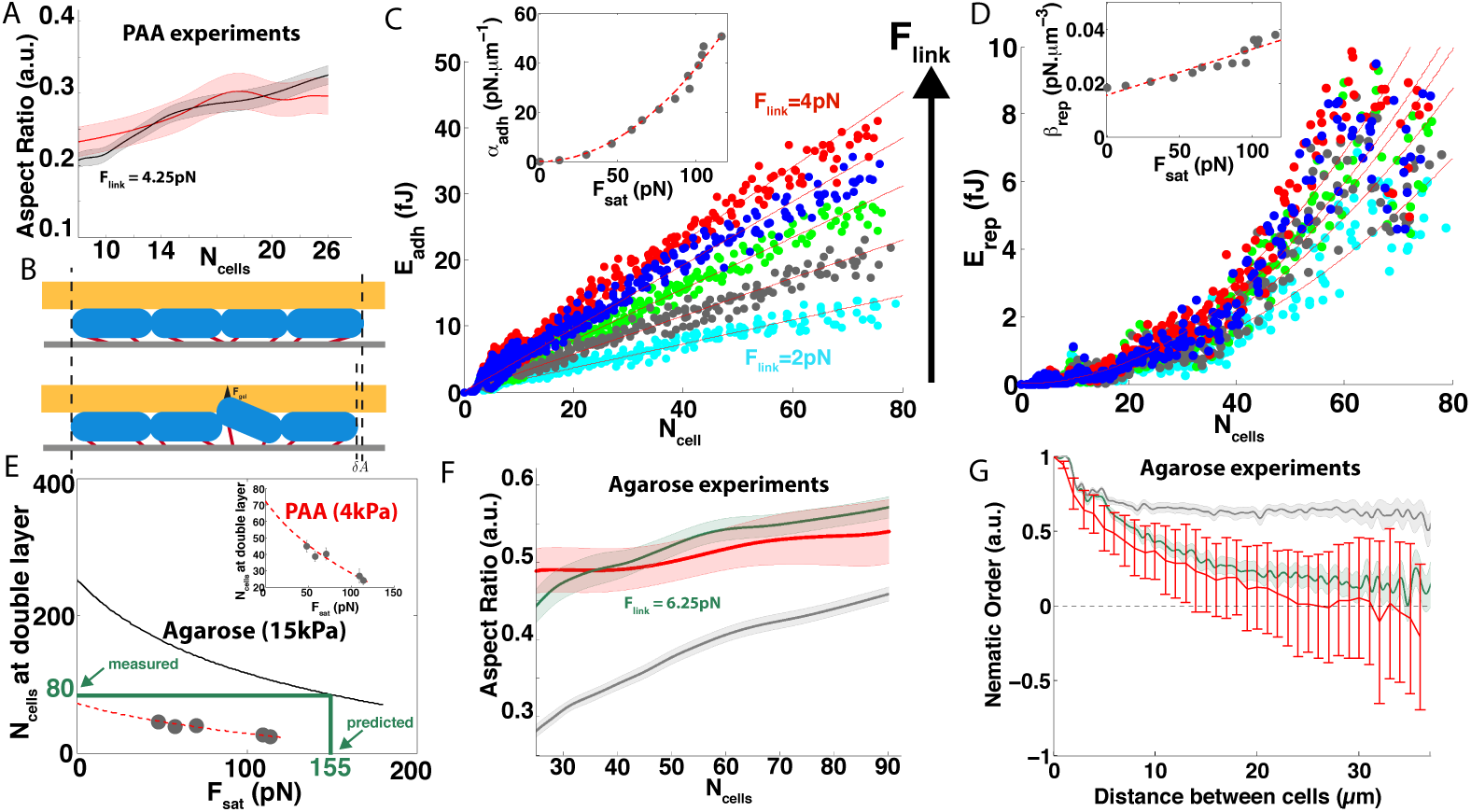
Simulation of microcolony morphogenesis.

A. Aspect ratio of WT *E. coli* microcolonies grown between agarose and PAA (red, N=26) compared to simulations (black; N=10, *F_link_* = 4.25*pN*). B. When the microcolony expands in the plane (upper panel), both the elastic energy *E_adh_* of the links and the repulsive energy *E_rep_* of steric interactions increases. On the contrary, when a bacterium expands in 3D it releases its repulsive energy and the microcolony saves a small surface element δA. Yet, the bacterium has to pay a cost in order to deform the gel and to extend its elastic links in the vertical direction. C. Numerical simulations shows that the adhesive energy stored in they increases with the number of bacteria in the microcolony *E_adh_* = *α_adh_ N_cells_*; the prefactoradh quadratically scales with the saturation force (inset). D. The repulsive energy between cells quadratically increases with the number of bacteria in the microcolony 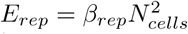; the prefactor *β_rep_* linearly increases with the saturation force (inset). In C and D, color codes for different values of *F_link_* at which adhesive links rupture (red, 4pN; blue, 3.5pN; green, 3pN; grey, 2.5pN; cyan, 2pN). E. Scaling between the size of the microcolony at double layer formation and the saturation force *F_sat_* (theory in red, experimental values reported in Fig. 3F in black). Increasing the stiffness of the gel, that the bacterium deforms, shifts the relationship (inset: red, PAA 4kPa; black, agarose 15kPa). F. From the value (horizontal green line) of the number of bacteria at the onset of the double layer for glass-agarose experiments (Fig. 1G), we inferred the saturation force *F_sat_* and the corresponding *F_link_* (vertical green line). F. Comparison between polar (green) and uniform (grey) simulations for the predicted *F_link_* and experimental data (red) for WT *E. coli* microcolonies grown between agarose and glass shown in Fig. 1F. G. The nematic order parameter 〈2cos^2^(θ_i,j_)-1〉_i,j_ as a function of the distance between two bacteria: experimental results for WT *E.coli* (red), simulation with polar adhesion (green) or uniform adhesion (grey) for the predicted *F_link_*. In F and G the simulations are defined as polar when adhesive links form only at the poles and uniform when they form all along the bacterium.

In summary, we experimentally showed that adhesion is polar and mature over one generation time. We theoretically demonstrated that asymmetric adhesion governs colony morphogenesis. The dynamics of maturation offers an efficient way for the colony to spread on a surface while maintaining a dynamic attachment with it. On the other hand, the polar localization of adhesion allow bacteria to rotate in order to relax the stress to which they are exposed. In a flow, rotation enables bacteria to align with stream lines in order to reduce the viscous drag exerted by the fluid [39, 40]. In our configuration, this degree of freedom leads to the formation of more circular microcolonies, which favors cell-cell interactions.

Two non-exclusive scenarios could account for polar adhesion at the scale of single bacteria. A first possibility is that adhesive factors are themselves asymmetrically distributed on the bacterial envelope, as we observed for Ag43. A second possibility is that the efficiency of adhesion is not uniform along the cell axis. The spatial dynamics of adhesive factors on the bacterial envelope follows from a competition between the expansion rate of the cell wall and the exposition rate of adhesive molecules on the surface, which itself results from synthesis and transport across the peptidoglycan. Thus, local modifications in the balance between those two terms would create inhomogeneities in the concentration profile along bacteria. Although cell wall synthesis is homogeneous along the cell axis during the first phase of cell cycle, cell wall expansion mostly occurs at the septum during the second phase [23]. This process could give rise to new poles with a lower concentration of adhesive factors. Similarly, it has been shown that virulence factors are distributed along a gradient from pole to pole, the slope of which is steeper at higher elongation rate [41]. Alternatively or concomitantly, bacterial adhesion could also have asymmetric efficiencies at the new and the old pole due to the elongation forces of the bacterium itself. If one pole is more anchored, it will remain fixed, promoting the formation of new links, while adhesive links at the other pole will more likely detach as they are put under tension during the cell cycle.

Finally, the shape of the microcolony depends on the overall level of cell-substrate adhesion and its subcellular localization. Uniform or lower levels of cell-substrate adhesion give rise to more elongated microcolonies, and thus more bacteria exposed to the environment at the boundary. In contrast, polar adhesion leads to more circular microcolonies and thus promotes cell-cell interactions inside microcolonies. Such a strategy could be particularly useful when facing environmental challenges. For instance, it has been shown that *P. aeruginosa*, which benefits from cell-cell contacts for siderophore usage, forms a second layer at smaller microcolony area when iron gets rare [19]. This indicates that adhesion may be up-regulated in response to the stress. In conclusion, the level of adhesion appears to be a means bacteria can use to tune the balance between cell-cell and cell-environment interactions. This could also be important with regard to evolution and survival strategies: in a chain of bacteria, cells experience the same environment and selection processes at the individual level dominate, whereas compact microcolonies generate spatial heterogeneities that could be more favorable for group selection.

## Authors contribution

MCD performed and analyzed the data from force microscopy, laser ablation and colony growth experiments. MA and MCD analyzed cell movements in microcolonies. ND set up the laser ablation experiments and did the asymmetric assays on single cells. CB did the immunofluorescence experiments on Ag43. TM, DB and ND did the modeling. TM and ND designed and wrote the code for the simulations. MB designed and coordinated force microscopy experiments. JMG and CB determined the set of *E. coli* strains and the relevant experimental conditions for *E. coli*. SL did so for *P. aeruginosa*. ND, SL and CQ co-advised MCD during her PhD. CB, JMG, SL and ND wrote the paper.

## Acknowledgments

We thank José Quintas Da-Silva and Carlos Gonzales for the design and construction of the mechanical parts. We thank Irene Wang for her help with force microscopy calculations. We thank Michael Elowitz for providing us with Schnitzcell, which was used to retrieve the lineage of the microcolonies. This work was supported by the Agence Nationale pour la Recherche ANR-12-JSV5-0007-01 (to MB), ANR-10-LABX-62-IBEID (to CB and JMG) and ANR-11-JSV5-005-01 (to ND).

## Supplemental Information

### Bacterial strains

Strains are described in Table S1. All *E. coli* strains are derived from strain MG1655 (E. coli genetic stock center CGSC#6300) and were constructed by λ red linear DNA gene inactivation method using the pKOBEG plasmid [42, 43] followed by P1vir transduction into a fresh *E. coli* background or alternatively by P1vir transduction of previously constructed and characterized mutation or insertion. We targeted the 4 major cell surface appendages of *E. coli*, i.e. flagella, type 1 fimbriae, Ag43 and curli, and the 4 known exopolysaccharides of *E. coli*, i.e. Yjb, cellulose, PGA and colanic acid. For *P. aeruginosa*, reference strain PA14 was used, as well as its fimbriae-deficient mutant cupA1:: MrT7, obtained from the PA14 transposon insertion mutant library (Ausubel lab, [44]).

### Microscopy and image analysis

Strains were inoculated in Lysogeny Broth (LB) from glycerol stocks and shaken overnight at 37°C. The next day, the cultures were diluted and seeded on a gel pad (1% agarose in LB). The preparation was sealed to a glass coverslip with double sided tape (Gene Frame, Fischer Scientific). A duct was previously cut through the center of the pad to allow for oxygen diffusion into the gel. Temperature was maintained at 34°C or 28°C using a custom-made temperature controller [45]. Bacteria were imaged on a custom microscope using a 100X/NA 1.4 objective (Apo-ph3, Olympus) and an Orca-Flash4.0 CMOS camera (Hamamatsu). Image acquisition and microscope control were actuated with a LabView interface (National Instruments). Segmentation [46] and cell lineage were computed using a MatLab code developed in the Elowitz’s lab (Caltech) [47]. For microcolony analysis, the cultures were diluted 10^4^ times in order to get a single bacterium in the field of view. Typically, we monitored 4 different locations and images were taken every 3 minutes in correlation mode [19].

### Morphological measurements

Experiments were performed on a confocal microscope (Leica, SP8). The aspect ratio is defined as a/b where a and b are respectively the large and the small characteristic sizes of the microcolony. We measured a and b by fitting the mask of the microcolony with an ellipse having the same normalized second central moments. The aspect ratio accounts for the anisotropy of the shape. It is close to zero for a linear chain of bacteria and close to one for a circular microcolony. All measurements are performed before the appearance of the second layer, which is detected manually in the time lapse sequence.

### Force microscopy

For experiments carried out at 34°C, bacteria were grown overnight in LB (37°C, 200 rpm), diluted 100-fold in 5 mL of fresh LB and grown again in the same conditions for 2 hours prior to experiments. Then the fresh culture was diluted 500 times and seeded on a 2% LB-agarose pad. To promote the expression of the curli fibers, some experiments were also carried out at 28°C and bacteria were taken from cultures at saturation. In that case, a 10^4^-fold diluted solution from an overnight culture at 28°C was directly seeded on the 2% LB-agarose-pad. In both conditions, a 4 kPa polyacrylamide (PAA) gel with 200nm fluorescent beads (FC02F, Bangs Laboratories) embedded below the surface was prepared [34]. The PAA gel bound to a glass coverslip was sealed onto the agarose pad with double-sided tape. Imaging was performed through the glass coverslip and the PAA gel with an inverted Olympus IX81 microscope. Fluorescence excitation was achieved with a mercury vapor light source (EXFO X-Cite 120Q). Beads were imaged through a 100X/NA 1.35 objective (Apo-ph3, Olympus) and an Orca-R^2^ CCD camera (Hamamatsu) with a YFP filter set (Semrock). The microscope, the camera and the stage were actuated with a LabView interface (National Instruments). Bacteria were imaged using phase microscopy. Force calculations were performed as previously described [48, 34]. A Fourier transform traction cytometry (FTTC) algorithm, with 0th-order regularization, was used to calculate the stress map from the substrate deformation, measured *via* the displacements of fluorescent beads embedded in the gel.

After correction for experimental drift, fluorescent beads were tracked to obtain a displacement field with high spatial resolution. The first frame of the movie, taken after seeding the sample with bacteria, was taken as the reference for non-deformed gel. The displacement field has been measured by a combination of Particle Imaging Velocimetry (PIV) and Single Particle Tracking (SPT). PIV was used to make a first measurement of the displacement field induced by the mechanical interactions between the micro-colony and the micro-environment. The obtained displacements where then applied to the reference bead image obtained for non-deformed gel. The relative displacement between this ?PIV-corrected? image and the deformed image was then analyzed using SPT to measure the residual displacement with a sub-pixel accuracy. The final displacement field was interpolated on a lattice of characteristic size 510 nm. Stress reconstruction was conducted with the assumption that the substrate was a linear elastic half-space medium. We set the regularization parameter to 10^−9^. To estimate the noise in stress reconstruction, we compared the average stress outside the colony where no forces are physically exerted, to the average stress beneath the microcolony (Figure S2). We derived the force *F_colo_* exerted by the microcolony on the substrate by integrating the mechanical stress over the surface covered by the microcolony. We then quantified the average stress *σ_colo_* beneath the microcolony by fitting the linear relation between *F_colo_* and the change in microcolony area. The maximal force was then simply obtained by multiplying the maximal value of the stress on the grid by the lattice elementary size (510 nm × 510 nm). To measure the asymmetry in force at the scale of the microcolony, we compared the maximal forces at new and old poles rather than the average forces because most of the pole slide on the substrate. The asymmetry of the microcolony is thus defined as follows: 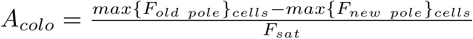.

### Single cell assays

**Asymmetric adhesion assays** Overnight cultures were diluted 10^2^ times in order to get on average 150 bacteria over 10 different fields of view; images were taken every 3 minutes in phase contrast. Because the two poles of a bacterium are not equivalent regarding their history [20], we project the displacement of the center of mass ∆*X* along the cell axis oriented towards the pole formed after the last division, i.e. the new pole. Since we do not know pole history until a division has occurred, we measured the absolute value of the parameter of asymmetry ∣*A*_cell_∣ in population of isolated cells. Then, we quantified the absolute value of the average 〈*A_cell_*〉 | by fitting the cumulative distribution of ∣*A*_cell_∣ with a folded normal distribution (Figure S3B).

**Single cell ablation** Ablations were performed using a UV pulsed laser (Explorer 349nm, Spectra Physics). A train of 30 impulsions at 1 kHz was sent, each delivering on the sample a power density of about 35 kW.µm^−2^. Using a custom algorithm on correlation images (Supp info), live image analysis enabled to automatically position the laser spot on a chosen bacterium by moving the stage (Thorlabs, MLS203). The setup is interfaced *via* LabView. These experiments were carried out on wild type *E. coli*, which larger size compared to *P. aeruginosa* enables successive ablations without perturbing the remaining cell (Figure S4B). Since we could not distinguish poles until one division had occurred, we computed the absolute value ∣*A_cell_*∣ for the first generation.

**Reorganization after division** Overnight cultures were diluted 10^3^ times in order to get on average 2 to 3 bacteria in each field of view; phase contrast images were taken in phase contrast every 30 seconds before septum formation and every seconds once the septum was visible.

**Immunostaining** Bacteria were grown to OD 0.2 and anti-ag43 was used at a dilution of 1:10,000. Immunostaining was performed in 1.5mL eppendorfs. Bacteria were then seeded between an agarose gel and a coverslip before image acquisition.

### Model for microcolony morphogenesis

Bacteria are modeled as spherocylinders that elongate exponentially at rate *g*, *d* = *gd*. They are allowed to divide at a constant rate once they have reached a size *d_L_*. Division occurs with a probability 1 if bacterial length exceeds 30% of *d_L_*. The dynamics of bacterial arrangement in the microcolony is driven by the two following effects: (i) cell-cell interactions are modeled with a Yukawa-like potential and (ii) cell-substrate adhesion are modeled by punctual elastic links that detach above a critical force. Adhesive links are created at the two poles with the same rate. In our model, asymmetric adhesion is a consequence of cell division that gives birth to new poles free of adhesive links. Once they detach, adhesive bonds are lost.

The interaction force between bacteria is derived from a Yukawa potential. For simplicity, we consider 6 balls (b) equally distributed along the cell’s center line. For each balls, we calculate interactions given the distance *r* between the points of distinct bacteria.

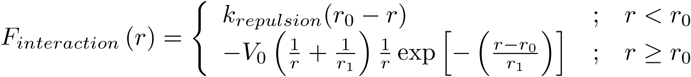

*F*_interaction_ is a force per unit length. *k*_repulsion_ is a constant. For each cell, the interaction terms are then multiplied by d/6, where *d* is the cell length. The distance of repulsion *r*_0_ sets the cell width. *V*_0_ sets the potential depth and *r*_1_ sets the range of attraction.

Adhesins or polysaccharides are polymers whose elasticity is derived from the Worm-Like-Chain model [49]:

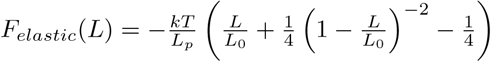

where *L*_0_ and *L_p_* are respectively the total and the persistence lengths of the polymer. *L* is the link extension.

Elastic links form at rate *k_on_* and saturates once they reached a number *n*_l_ per ball. In the polar case, they are randomly distributed at both poles in a disk of diameter *r*_0_. In the uniform case, they are randomly distributed all along the spherocylinder. They detach at a rate that depends on the tension exerted on the link:

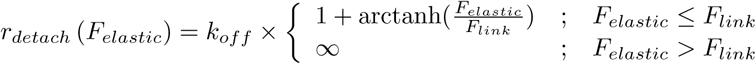

The threshold in force *F_link_* is a parameter used to vary the strength of adhesion.

We perform over-damped molecular dynamics simulations to model the motion of bacteria:

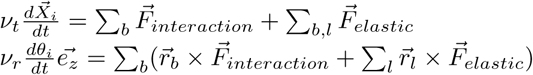

where *i*, *b* and *l* are respectively the indices for the cells, the balls that constitute a cell and the adhesive links between the cells and the substrate. *v_t_* and *v_r_* are the translational and rotational friction coefficients for cylinders in a viscous fluid of viscosity *η* [50]:

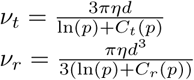

where 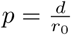 is the aspect ratio of bacteria. *C_t_* and *C_r_* are given by:

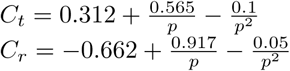

As bacteria elongate, adhesive links with the substrate are extended and surrounding bacteria are pushed. As a result, the elastic and the repulsive energies increase as the microcolony expands in 2D. We compute the elastic energy and the repulsive energy as follows:

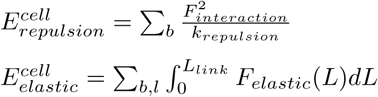

The scaling of the total energy in the microcolony deduced from our simulation is the following (Fig. 4C,D):

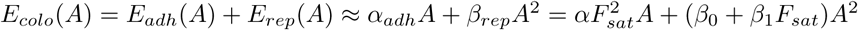

Bacteria proliferate in 2D until it is less energetically favorable for the microcolony to exclusively raise the energy in the plane rather than paying the cost required for a bacteria to deform the gel and go 3D. We compared the two situations with a mean-field argument:

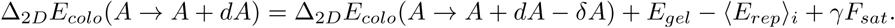

The left-hand side corresponds to a situation in which all the increase in surface area remains confined to the plane. The right-hand side of the equation corresponds to a situation in which a bacterium expands into the third dimension, thus saving a small surface extension δA. The bacteria that deforms the gel in 3D has to pay a deformation cost *E_gel_* and a mechanical work _γ_*F_sat_* to elongate its adhesive links in the third dimension. However, the cell that expands into the third dimension gains its repulsive energy *E_rep_*. The energy balance between the two situations can be rewritten as follows, where *A*_0_ is the average area of bacteria after division:

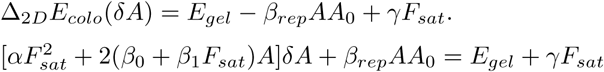

During this transition, the work of the adhesive links is proportional to the rupture force of individual links *F_link_*, itself proportional to *F_sat_* and to the elongation of the links *z* corresponding to the indentation of the gel. Thus _γ_*F_sat_* = _*γ*0*z*_*F_sat_* Similarly, we estimate that *δA* = *zr_0_*.

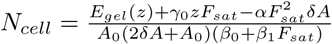

We used the Hertz model to compute the energy required to deform the softer interface:

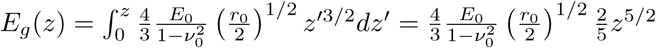

*E_0_* and *v_0_* are respectively the Young modulus and the Poisson ration of the soft interface.

The values of the parameters used in this study are referenced in Table S3.

### Movie S1

**WT *E. coli* microcolony grown between agarose and glass**. Phase contrast images were taken every 3min before septum formation. Time is displayed hours and minutes. Scale bar 10μm

### Movie S2

**Δ_4_adh *E. coli* microcolony grown between agarose and glass**. Phase contrast images were taken every 3min before septum formation. Time is displayed hours and minutes. Scale bar 10μm.

### Movie S3

**Asymmetric adhesion during the first cell cycle**. Phase contrast images were taken every 3min. The arrows depict the initial position of the cell. Time is displayed in minutes. Scale bar 5μm.

### Movie S4

**One-sided and two-sided ablations**. Correlation images were taken every 3min. After each division, an ablation is performed either on the same side as the previous one (one-sided mode, movie on the left) or on the opposite side to the previous one (two-sided mode, movie on the right). In the one-sided mode, the pole of the remaining daughter bacterium is aging. On the contrary, it is rejuvenating after each division in the two-sided mode. Red and blue dots respectively indicate new and old poles. Time is displayed in hours and minutes. Scale bar 3μm.

### Movie S5

**Reorganization after division**. Phase contrast images were taken every second after septum was visible. Time is displayed in minutes and seconds. Scale bar. 3μm.

### Movie S6

**Force microscopy**. Map of the stress that bacteria exert on the polyacrylamide gel. Images were taken every 3min. The color code span from blue to red corresponding to 0Pa and 200Pa respectively (check). Time is displayed in hours and minutes. Scale bar 5μm.

### Movie S7

**Simulation of microcolony growth**. Forces exerted on the substrate are colored in red.

### Movie S8

**Simulation of microcolony growth**. Bacteria are shown in grey, the density of adhesive links is labelled in green.

---

**Figure S1.**
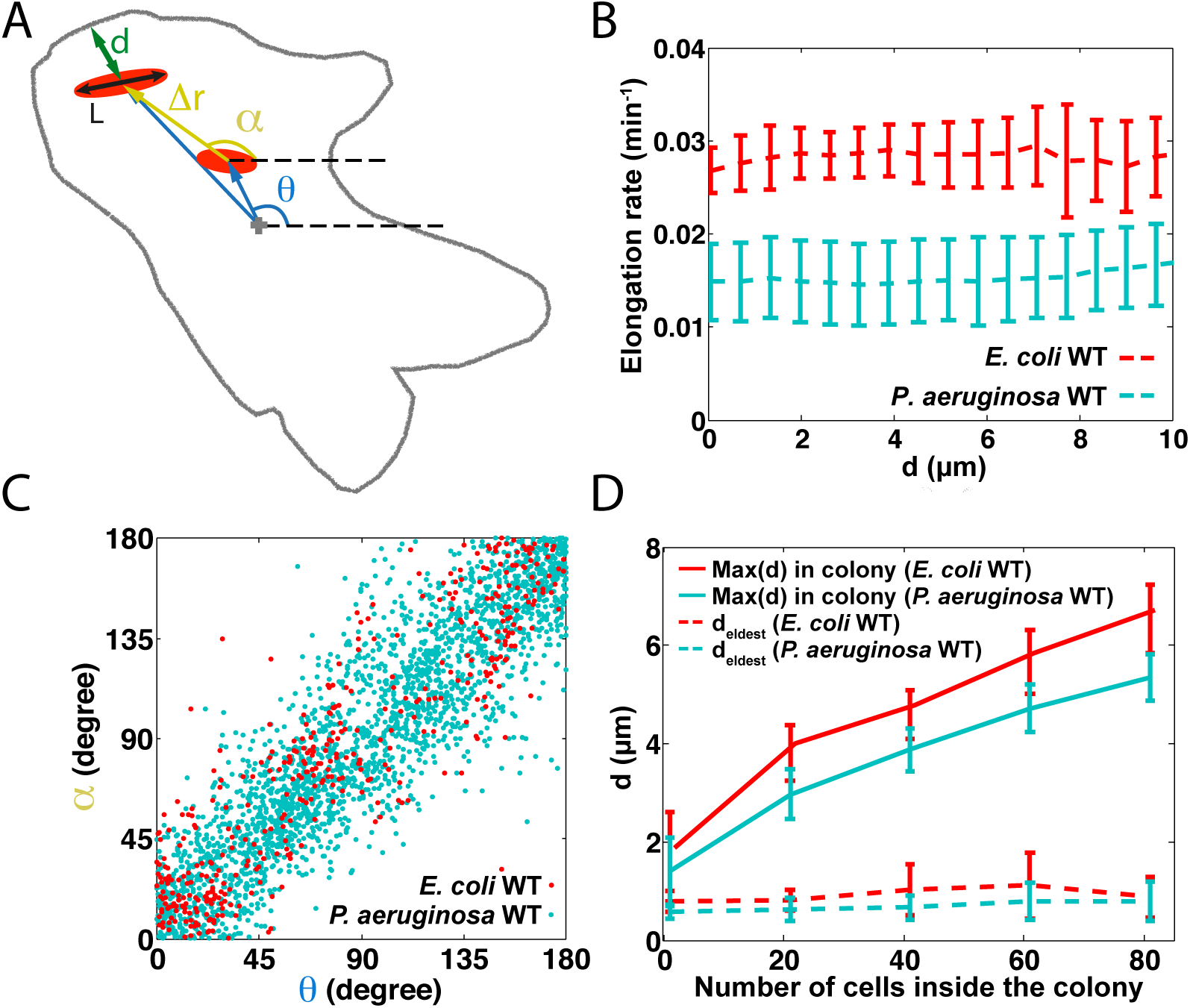
Bacterial elongation pushes the old cell to the periphery.

A. Parameters used to quantify the cell movements occurring during each cell cycle: cell length (L), average distance to the edge of the colony (d), angular position averaged over the cell cycle in the microcolony (*θ*), angular direction (*α*) and magnitude (Δ*r*) of the cell movement from the beginning to the end of the cell cycle. B. The elongation rate 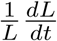 is uniform inside the microcolony. C. Correlation between the orientation of individual cell movements α and the angular position of *θ* bacteria. E. Distance of the oldest cell from the edge of the microcolony (dashed line) as a function of the number of cells in the microcolony. The solid line displays the characteristic size of the microcolony, computed as the maximal distance to the edge that can be measured for each cell inside the colony. In all panels, data are plotted in red for WT *E. coli* (N=7 microcolonies) and in cyan for WT *P. aeruginosa* PAO1 (N=10 microcolonies) and error bars represent standard deviations. The microcolonies were monitored up to the double layer formation.

**Figure S2.**
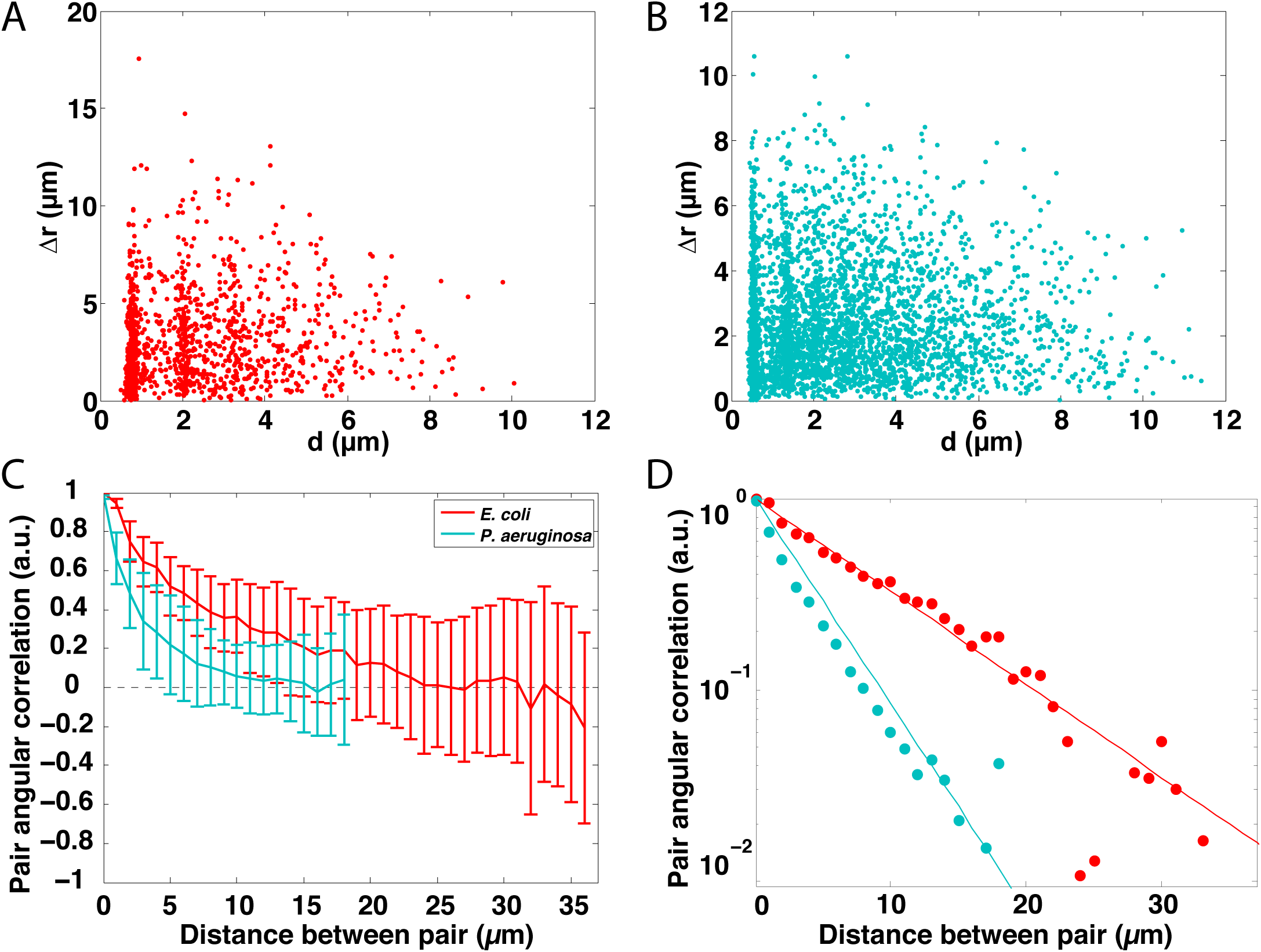
Bacterial orientation inside the microcolony.

A, B. The displacement Δ*r* of bacteria between their birth and their division as a function of their average distance d to the edge over the cell cycle. Each point represents a cell and data from different microcolonies are superimposed. A, WT *E.coli* (N=7 microcolonies, red); B, WT *P. aeruginosa* (N=10 microcolonies, cyan). C. The nematic order parameter in 2D 〈 2cos^2^(θ_*i*,*j*_) − 1〉_*i*,*j*_ as a function of the distance between two bacteria for microcolonies counting around 100 cells (*θ*_*i*,*j*_ is the angle between bacteria *i* and *j*). The same graph is plotted in log-lin scale in D. WT *E.coli* (N=7 microcolonies, red); Right, WT *P. aeruginosa* (N=10 microcolonies, cyan). Error bars represent standard deviations.

**Figure S3.**
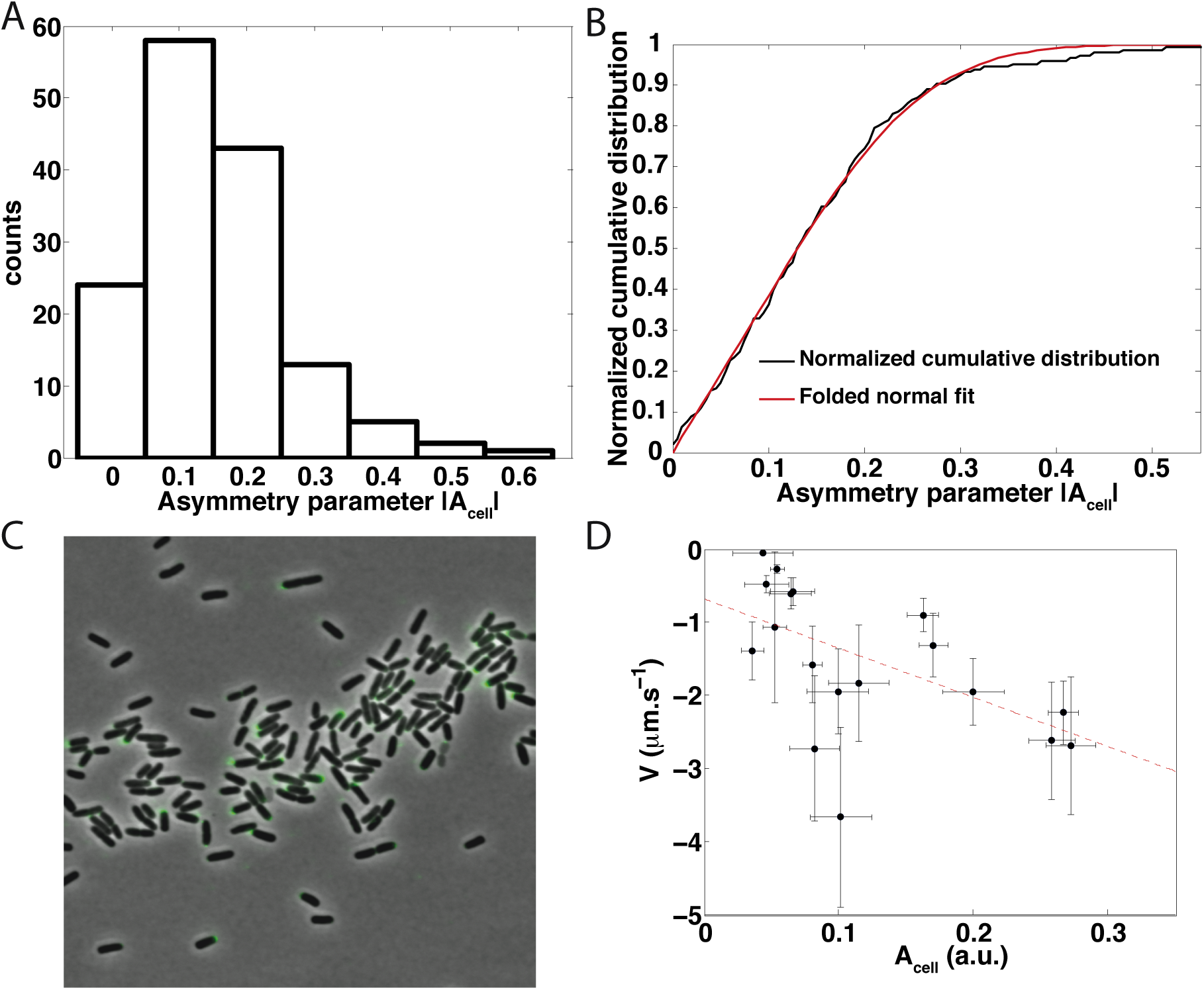
Asymmetric adhesion induces the reorganization of the two daughter cells after division.

A. Histogram of | *A_cell_* | for a wild type population of *E. coli* cells. B. Normalized cumulative distribution (black line) of the histogram shown in A is fitted with a normalized cumulative distribution a folded normal law (black line). C. Immunodetection of Ag43 in a WT *E. coli* culture started from a colony expressing Ag43, using an anti-Ag43 polyclonal rabbit antiserum raised against the α-domain of Ag43. Because the expression of ag43 is bistable [51], not all the bacteria are expressing Ag43. Yet, for the fraction of cells that express it, Ag43 is localized only at one pole. D. Magnitude of the reorganization of the two daughter cells as a function of the mother cell asymmetry *A_cell_*. The magnitude of the reorganization is computed as the maximal derivative of cell-cell distance measured during the 100 seconds after this distance reached its maximum. The sign of reorganization is negative since the two daughter cells are moving closer. For comparison, the sliding speed due to cell elongation is on the order of 5*nm.s*^−1^.

**Figure S4.**
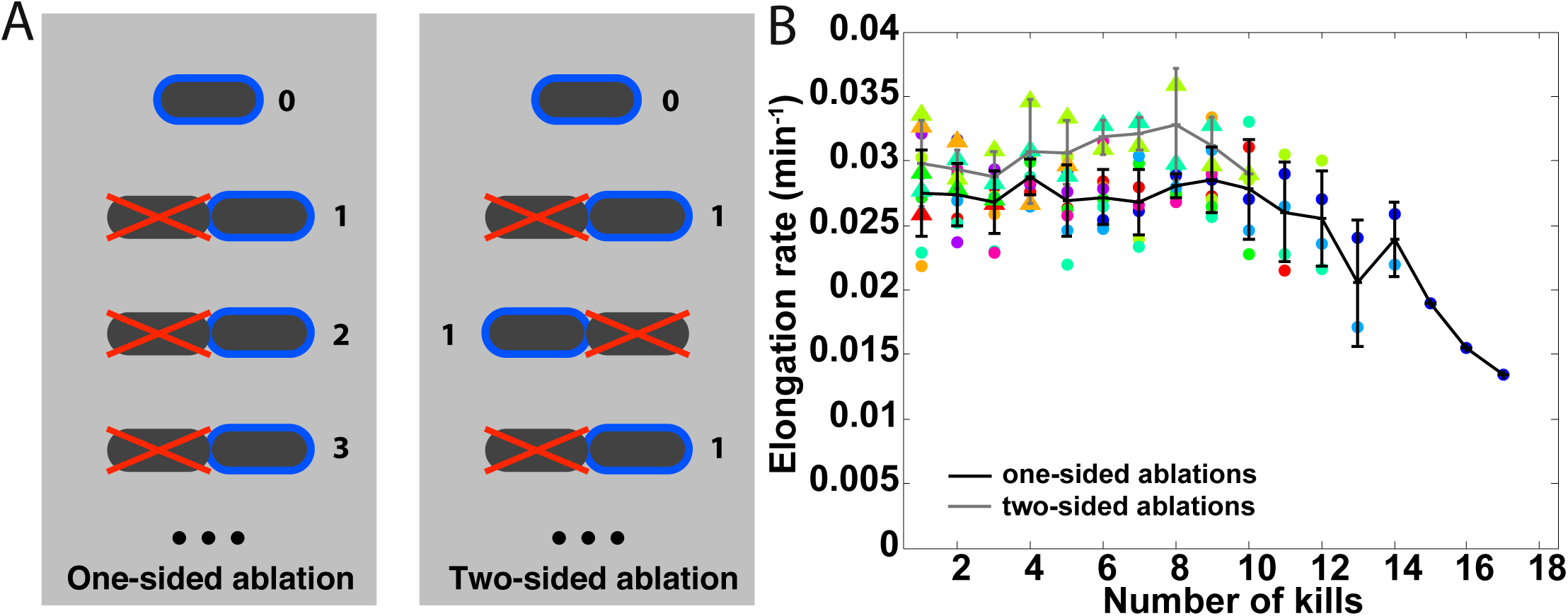
One-sided and two-sided ablations.

A. For one-sided ablations, we systematically ablated the left daughter cell after division. Therefore, the oldest pole of the remaining cell is aging throughout the successive rounds of ablations. For two-sided ablations, the ablations were alternated between the left and the right daughter cell at each generation. Therefore, the oldest pole of the remaining cell was never older than one generation throughout the successive rounds of ablations (except for the two first rounds). The numbers illustrate the age of the oldest pole, which starts arbitrarily at 0. The red cross indicates the ablated cell. Time in generations (i.e. one cell cycle) goes from top to bottom. B. The elongation rate of the remaining cell as a function of the number of cell cycles during the course of one-sided ablation (black line) and two-sided (gray line) experiments. Error bars represent the standard deviations. The different points represent individual experiments corresponding to one-sided (dots) and two-sided ablations (triangles).

**Figure S5.**
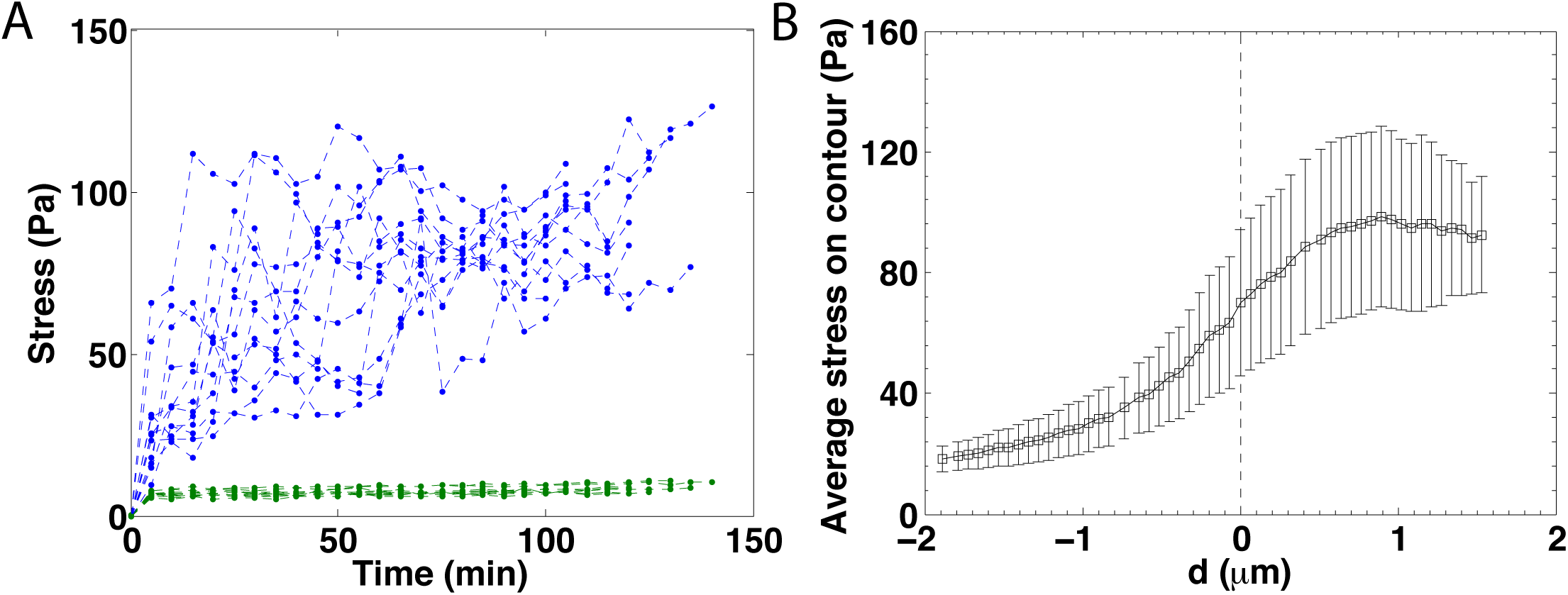
Stress beneath and outside of the microcolony..

A. Time evolution of the stress beneath (blue) and outside (green) microcolonies of WT *E. coli*. B. Average adhesive stress as a function of the distance d to the edge of WT *E. coli* microcolonies. Microcolonies are sliced in concentric contours going from the edge to the center. From the stress maps shown in Fig. 3, we measured the mean stress inside the contour computed as the integral of the stress along the contour divided by the length of the contour. The edge of the colony is set at *d* = 0. The inside of the microcolony corresponds to *d* > 0. Error bars represent standard deviations of the stress distribution in concentric contours.

**Figure S6.**
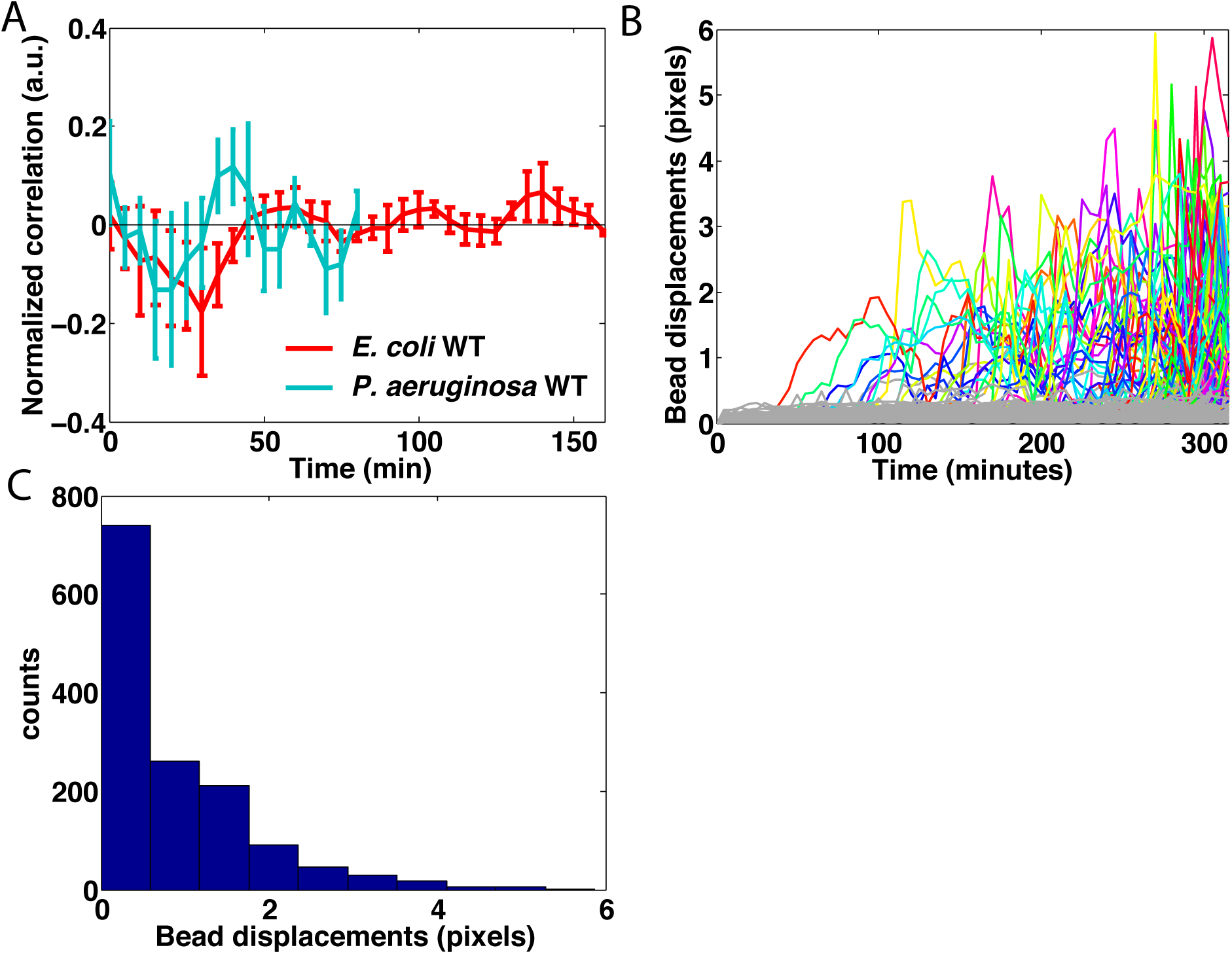
Rupture of adhesive bonds.

A. Average correlation between pole displacement and force variation. in wild type microcolonies of *E. coli* (red, N=12) and *P. aeruginosa* (cyan, N=16). Poles are tracked from their birth (beginning of pole history) and spans several generations (example of an individual trace is shown in Fig. 3G). B. Displacement of individual beads located in the PAA gel under the colony (colors) and outside the colony (gray). Time 0 refers to the start of the experiment. All individual trajectories from an individual microcolony growth are displayed. Displacements are computed as the distance between the position of the beads at a given time relative to their position on the first frame. C. Histogram of the bead displacements for beads located inside the PAA gel under the microcolony. Data of all time points shown in B are pooled in the distribution.

**Figure S7.**
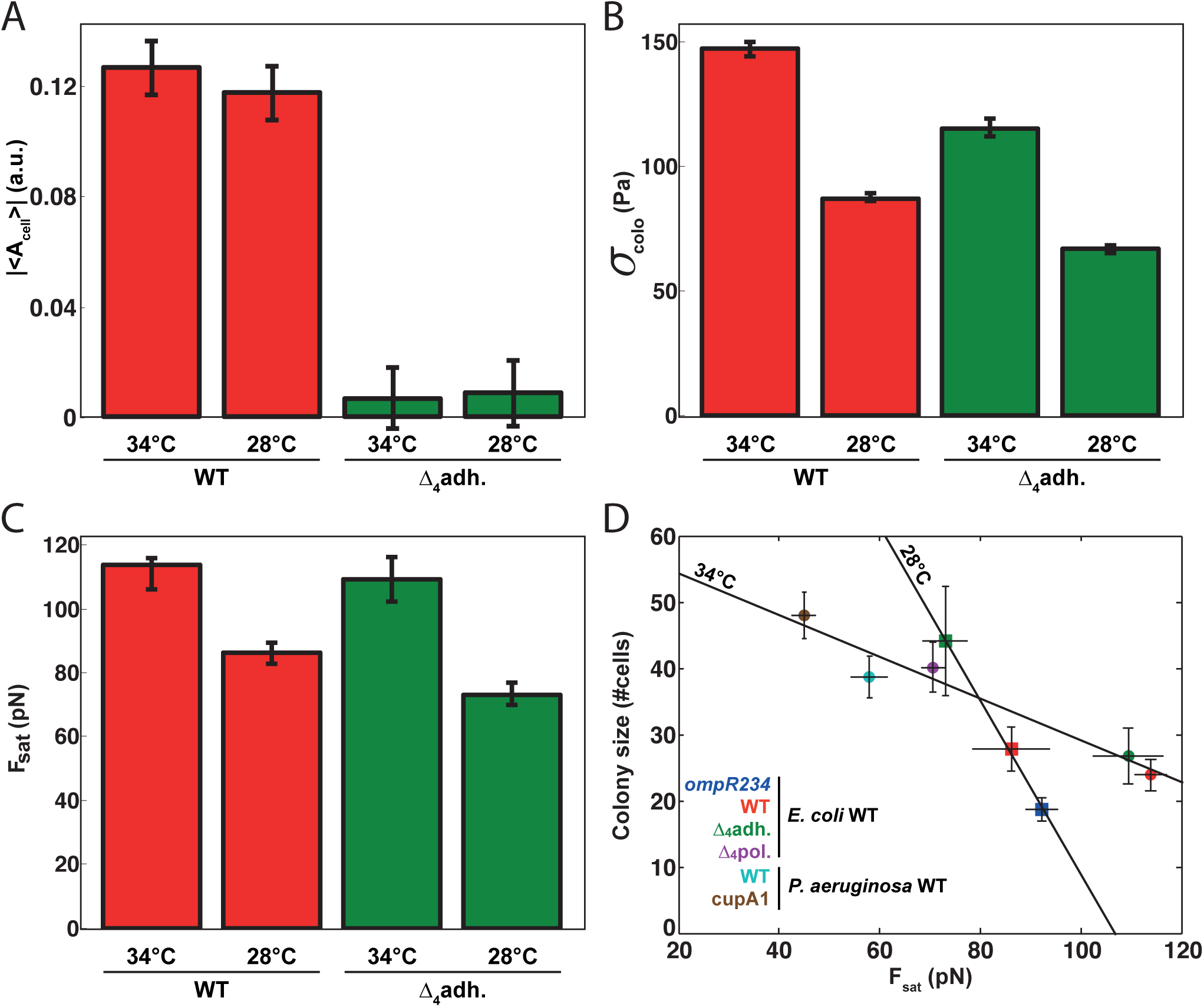
Direct comparison of adhesive properties at 28°C and 34°C.

A. Average asymmetry | 〈*A_cell_*〉 | for a population of isolated bacteria for WT *E. coli* (red, N=142 for 28°C, N=146 for 34°C) and a mutant deleted for four adhesins Δ_4_adh, Δ*fliER_agn43_fimAH_csgA* (green, N=115 for 28°C, N=95 for 34°C). *σ_colo_* (B) and *F_sat_* measured by force microscopy experiments for WT *E. coli* (red, N=13 for 28°C, N=12 for 34°C) and a mutant deleted for four adhesins Δ_4_adh, Δ*fliER_agn43_fimAH_csgA* (green, N=7 for 28°C, N=19 for 34°C). D. Number of bacteria at the onset of the second layer formation as a function of the saturation force for all strains considered in this study at 28°C and 34°C.

**Figure S8.**
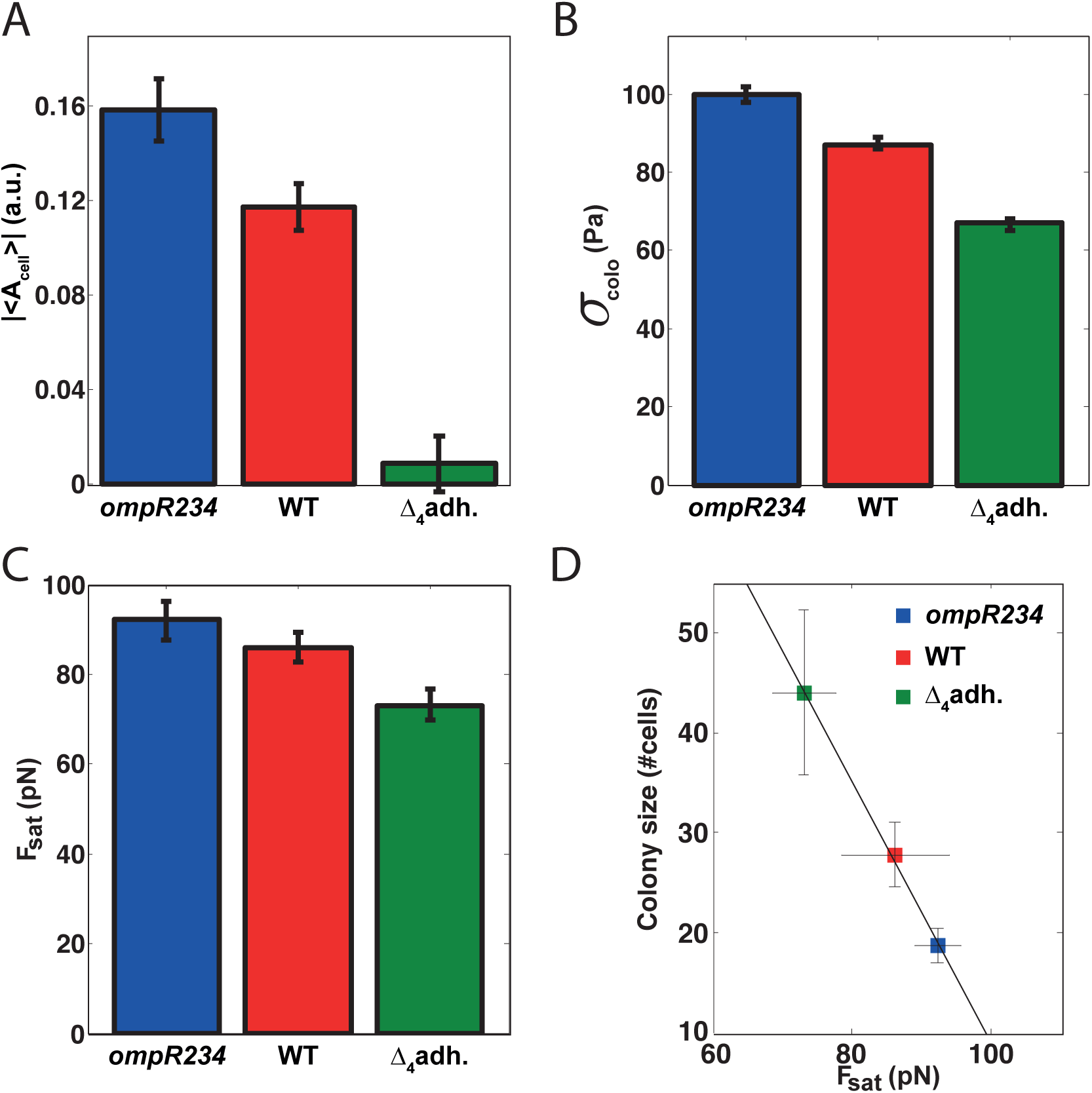
Cell-substrate adhesion at 28°C.

A. Average asymmetry |〈*A*_*cell*_〉| for populations of isolated bacteria for WT *E. coli* (red, N=142), for a mutant deleted for four adhesins Δ_4_adh, Δ*fliER_agn43_fimAH_csgA* (green, N=115) and for mutant *ompR234* (blue, N=82) which over-expresses curli fibers. *σ_colo_* (B), *F_sat_* (C) and number of bacteria at the onset of the second layer formation as a function of the saturation force (D) measured by force microscopy for WT *E. coli* (red, N=13), a mutant deleted for four adhesins Δ_4_adh, Δ*fliER_agn43_fimAH_csgA* (green, N=7) and a mutant *ompR234* (blue, N=10) which over-expresses curli fibers.

**Figure S9.**
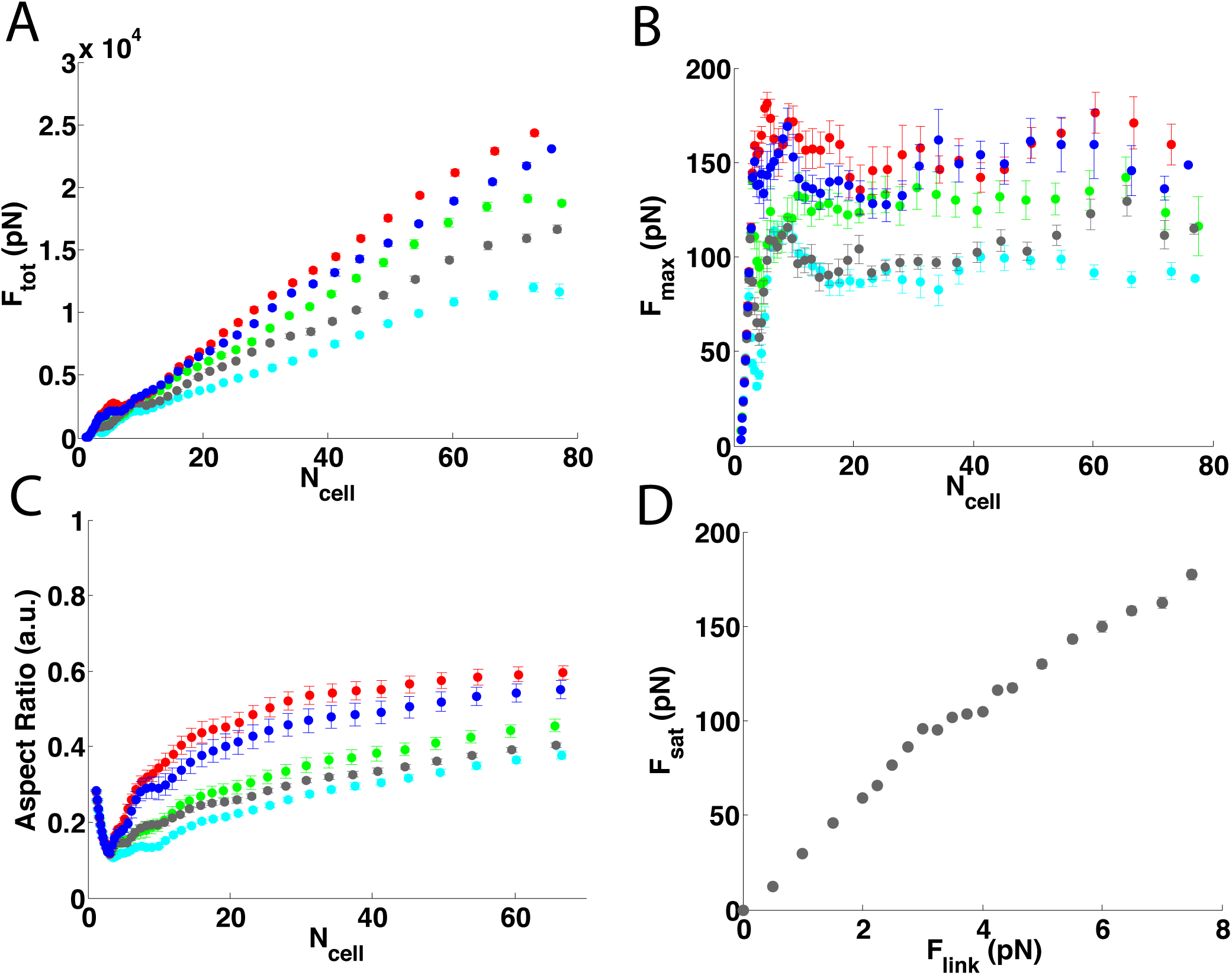
Simulation of microcolony morphogenesis.

A. *F*_*tot*_ as a function of the colony size. B. The maximal force *F*_*max*_ as a function of the colony size. We used these curves to fit the value of the plateau *F*_*sat*_ for the simulations. C. Aspect ratio for different microcolonies. Each curve is the average of 10 simulations corresponding to *E* coli size for different values of the detachment force for individual links *F*_*link*_: 2*pN* (cyan); 2.5*pN* (grey); 3*pN* (green); 3.5*pN* (blue); 4*pN* (red). D. Relation between the saturation force *F_sat_* and the rupture force *F_link_* of single adhesive links.

**Figure S10.**
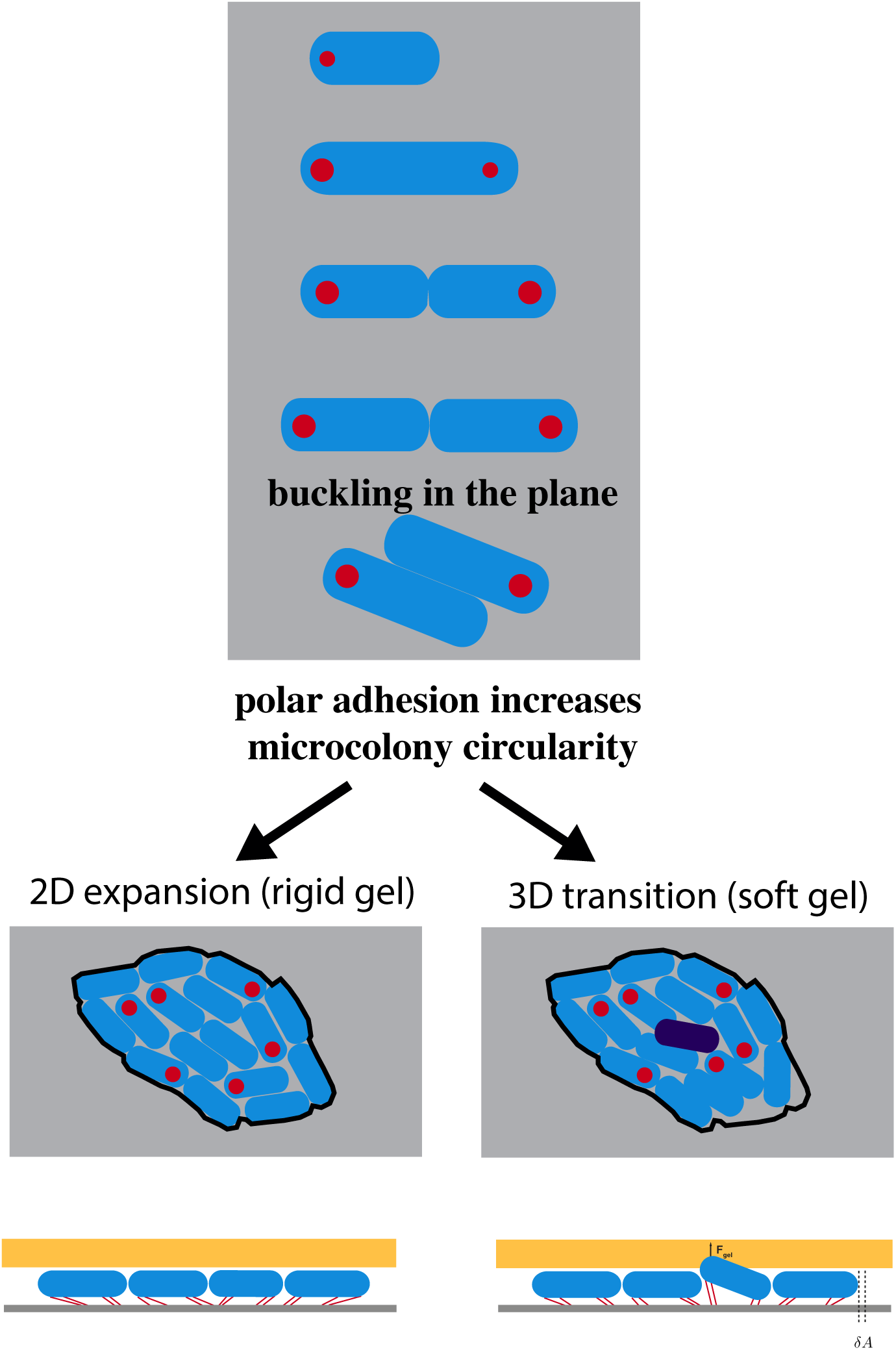
Schematic of microcolony morphogenesis.

Polar adhesion forces daughter cells to slide along each others and subsequently increases the circularity of the microcolony. On soft gel, the microcolonies form double layers at smaller sizes than on rigid gel. The black boundary depicted in the two cases is the same. Red points are used to schematize polar adhesion. The cell highlighted in purple is the cell that starts to grow on top of the others.

**Table S1** Strain description.

**Table 1.**
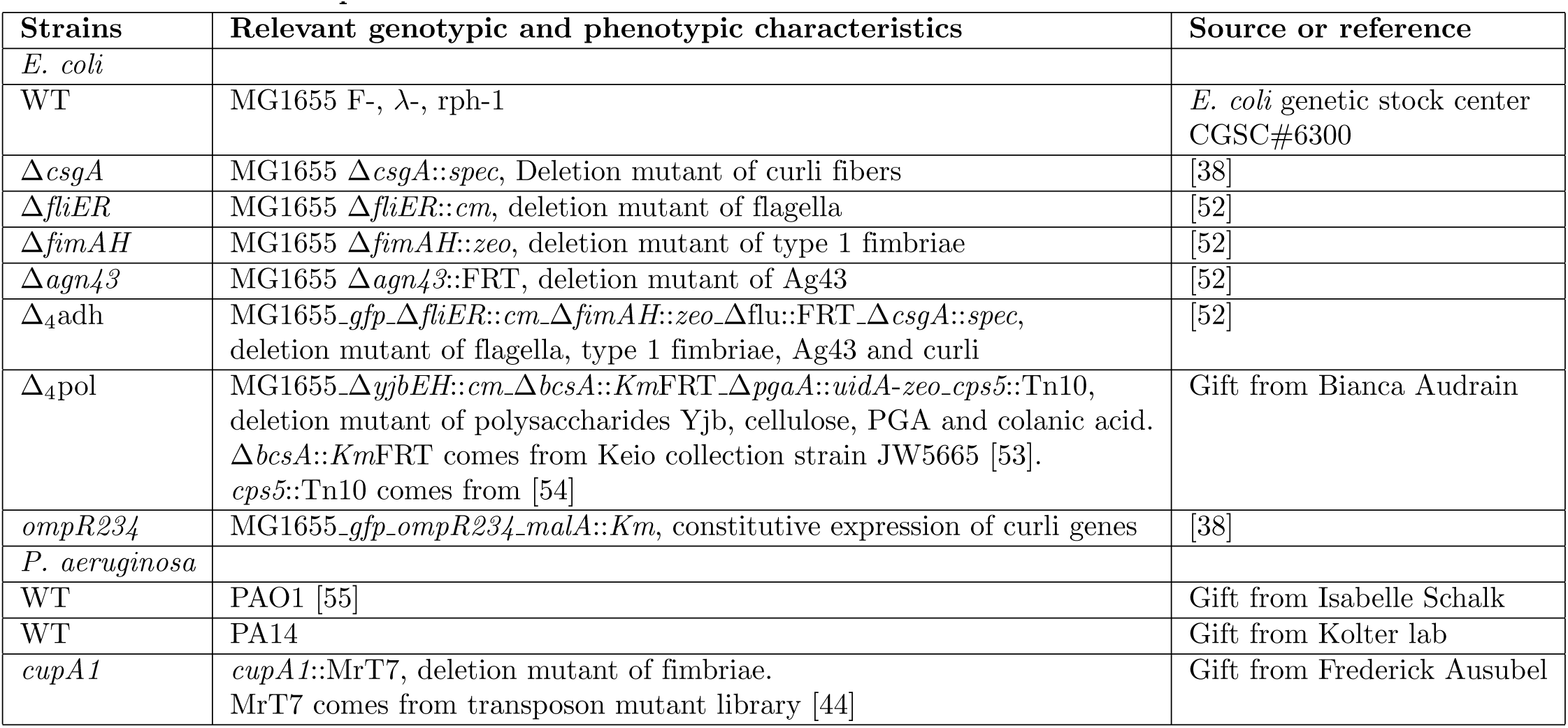
Strain Descritpion.

**Table S2** Primer description.

**Table 2.**
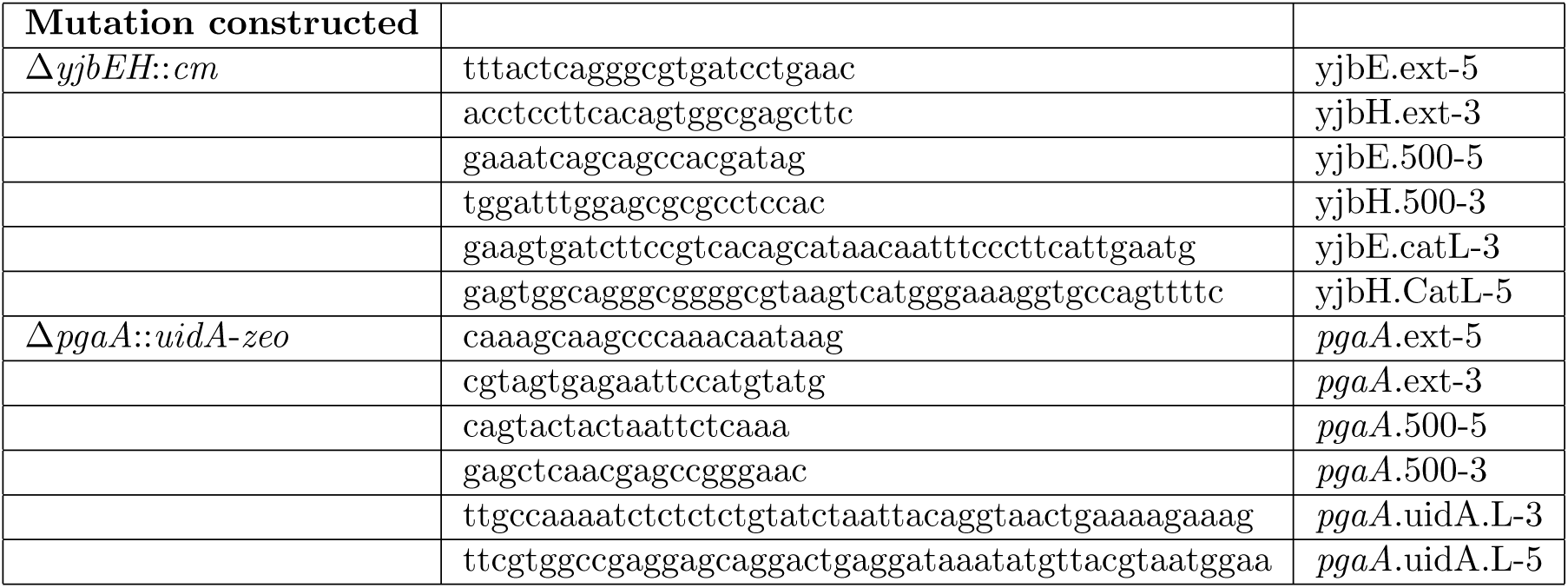
Primers used in this study to construct deletion mutants.

**Table S3** Values of the parameters involved in the simulation.

**Table 3.** Value of the different parameters described in the model.

| Name | Value | Description | Reference |
| --- | --- | --- | --- |
| $dt$ | 2s | time increment | |
| $g$ | $1.6h^{-1}$ | growth rate | *This study* |
| $r_0$ | $1.4\mu m$ (*coli*); $0.9\mu m$ (*aeruginosa*) | cell width | *This study* |
| $d_L$ | $8.8\mu m$ (*coli*); $4.6\mu m$ (*aeruginosa*) | cell length at which cell may divide | *This study* |
| $k_{repulsion}$ | $10^4 pN/\mu m$ | elastic constant for cell repulsion | [52] |
| $V_0$ | $2.5pN.\mu m$ | depth of the attractive potential | |
| $r_1$ | $0.1\mu m$ | attraction length | |
| $k_{on}$ | $22.5min^{-1}$ | rate of adhesive link formation per cell | |
| $k_{off}$ | $10^{-2}min^{-1}$ | rate of adhesive link loss | |
| $n_l$ | 250 | maximum links per cell | |
| $L_p$ | 0.828nm | persistence length | [56] |
| $L_0$ | $10\mu m$ | length of a fully extended link | [36] |
| $F_{link}$ | few pN | force at which single adhesive links break | [57] |
| $\eta$ | $2pN.s/\mu m^2$ | coefficient of the viscous friction with the walls | |
| $E_0$ | 4kPa | Young modulus of the above gel | *This study* |
| $\nu_0$ | 0.5 | Poisson ratio of the above gel | |
| $z_0$ | $0.8\mu m$ | Vertical indentation of the gel at double layer formation | |

